# Quantifying RNA Folding Equilibria from Unclassified Cryo-EM Particle Images

**DOI:** 10.64898/2026.10.11.758367

**Authors:** Debanjana Maji, Lars Dingeldein, Minhuan Li, Miro A. Astore, Luke Evans, Wai Shing Tang, Jutta Rogal, Aaditya Rangan, Roberto Covino, Alex de Marco, Steve L. Bonilla, Pilar Cossio

## Abstract

Functional RNA structures are typically far more dynamic than most proteins solved by cryo-electron microscopy (cryo-EM). Flexible junctions and weak tertiary contacts produce broad conformational ensembles, and cryo-EM particle classification analyses discard unaligned particles, which inherently contain the information needed to quantify such broad ensembles. Here, we infer compositional and conformational distributions from unclassified particle images by Bayesian ensemble reweighting. We tested our approach on the P4-P6 domain of the *Tetrahymena thermophila* group I intron, a classic model system of Mg²⁺-dependent RNA folding. While stable monomeric and dimeric states yielded high-resolution reconstructions, analyses of abundant, flexible conformations required Bayesian ensemble inference. Our approach quantitatively captured the Mg²⁺-dependent folding transition, indicating that cryo-EM can report the abundance of coexisting RNA states in agreement with bulk solution measurements. These results showcase how quantitative studies of RNA folding can be achieved by combining cryo-EM measurements with statistical ensemble-inference tools.

**SUMMARY:** RNAs are remarkably versatile macromolecules that fold into dynamic three-dimensional (3D) structures to perform critical functions in cells and viruses, making them increasingly attractive target for therapeutics and as tools for biotechnology. Yet our understanding of RNA structure–function relationships remains limited, in part because structural biology tools are poorly suited to probing the dynamic ensembles that RNAs occupy. The raw 2D images collected in single-particle cryo-EM, however, inherently contain the information needed to visualize and quantify such ensembles. Here, we use the P4-P6 domain of the *Tetrahymena thermophila* group I intron, an archetypal model of Mg²⁺-dependent RNA folding, to leverage this information. While stable monomeric and dimeric states yielded high-resolution structures, flexible conformations escaped conventional 3D reconstructions—even though they dominate at physiological Mg²⁺ concentrations. To overcome this limitation, we integrated single-particle cryo-EM with a Bayesian reweighting framework that quantifies RNA ensemble distributions directly from raw 2D images. By statistically reweighting computational ensembles against experimental images via probabilistic modeling and deep learning, our approach quantitatively captures Mg²⁺-dependent population shifts between extended, compact, monomeric, and dimeric species, in agreement with fundamental principles of RNA folding and bulk solution measurements. Our framework supports the sampling and visualization of conformations within the ensemble, allows the quantification of errors, and demonstrates that the learned continuous ensemble fits the data significantly better than any set of discrete structures. These results showcase how quantitative RNA folding can be achieved by combining high-resolution structural biology measurements with statistical tools that treat RNAs as conformational ensembles.

## INTRODUCTION

Beyond encoding genetic information, RNAs fold into diverse three-dimensional (3D) structures that carry out critical functions in cells and viruses (*1, 2*). Well-established examples include tRNAs, group I and II introns, small self-cleaving ribozymes, and riboswitches, among many others (*3–7*). Given their essential biological role, structured RNAs are increasingly recognized as promising therapeutic targets and tools for biotechnology, acting as aptamers, biosensors, and programmable nanostructures (*8–11*). However, our current understanding is limited by the dynamic nature of RNA structure. Folded RNAs exist as dynamic conformational ensembles rather than as single, well-defined states, making them difficult to characterize by conventional structural biology tools (*1, 12–14*). Despite this challenge, two properties of RNAs make the understanding of such ensembles more tractable than it might otherwise be: hierarchical folding and structural modularity.

Structured RNAs fold through secondary structure base pairing, and the resulting helices are subsequently organized in 3D space through tertiary folding (Supplementary Figure 1) (*15–17*). This organization is guided by the conformational preferences of flexible junctions and by the formation of sparsely distributed tertiary interactions (*18–20*). A practical consequence of this hierarchy is that secondary-structure formation and tertiary folding can often be probed independently of each other (*16, 17*). In contrast to the highly cooperative folding characteristic of globular proteins, the hierarchical nature of RNA folding leads to modular architectures, with independently folded secondary structural elements that can form in isolation (*18*). Together, hierarchical folding and structural modularity offer a modeling framework for dissecting RNA folding. Yet the central difficulty remains – flexible junctions and weak tertiary interactions leave RNAs sampling broad conformational ensembles that traditional structural biology techniques are poorly suited to capture, since most techniques rely on averaging.

Due to its ability to capture snapshots of individual particles and resolve conformational heterogeneity through particle classification, cryo-electron microscopy (cryo-EM) has emerged as a powerful technique for visualizing RNA structures (*21–26*). In principle, the 2D particle images contain the information needed to quantify conformational ensembles (*27, 28*), including those of RNA (*12*). However, traditional single-particle cryo-EM 3D reconstruction remains limited when dealing with highly dynamic RNAs, and whether Boltzmann-distributed ensembles can be recovered from cryo-EM data remains an open question (*29, 30*). Typical cryo-EM workflows depend on aligning each particle to a discrete class, whereas RNA structures tend to populate broad ensembles that challenge alignment and discrete classification (*12, 24, 31*). Although methods have been developed to model heterogeneity in cryo-EM using manifold embedding, dimensionality reduction, and deep learning, most rely on pose alignments determined from a consensus reconstruction (*28, 32–36*). These approaches, therefore, describe conformational flexibility near a well-defined state but fail to capture conformations that deviate substantially from it (*28, 37*). However, a mechanistic understanding of RNA structure-function relationships requires a full description of the ensemble (*1*). This information is present in the raw single-particle images, but it is lost in conventional analyses, as the particles that resist alignment into consensus maps are discarded.

To address these challenges, we apply a Bayesian reweighting framework to dissect 3D conformational and compositional distributions of RNAs, and to quantitatively assess how these distributions depend on ionic conditions and mutations. Using the P4-P6 domain of the *Tetrahymena thermophila* group I intron as a model system, we first solve cryo-EM reconstructions of the major states populated across different Mg^2+^ concentrations ([Mg^2+^]) and find that a large fraction of particles—predominant at physiological [Mg^2+^]—cannot be resolved. We then use the reconstructions as starting points for computational modeling of broad 3D ensembles, leveraging hierarchical folding and structural modularity to make the conformational search tractable. The computational ensembles serve as a prior for Bayesian reweighting, in which we infer the relative populations of compositions and conformations using a simulation-based inference likelihood evaluated on raw single-particle cryo-EM projection images. Our quantitative analysis of P4-P6 across [Mg^2+^] is consistent with solution experiments but reveals a much richer conformational and compositional landscape than previously described for this model RNA. Our work establishes cryo-EM as a quantitative tool for measuring RNA ensembles and their dependence on sequence and environmental factors, with implications to the study of RNA folding, self-assembly, and ligand recognition under biologically relevant conditions.

## RESULTS

### A model system for the study of RNA conformational ensembles

Study of the 55 kilodalton P4-P6 domain of the *Tetrahymena thermophila* group I intron ribozyme has provided fundamental insights into RNA folding, electrostatics, and structural modularity (*19, 38–46*). Its conformational landscape has been primarily described in terms of two states: open and closed (Figure 1A) (*19, 40*). The closed state was previously visualized by X-ray crystallography, where P4-P6 crystallized as a dimer (Figure 1B) (*38*). It contains several 3D motifs that are ubiquitous across folded RNAs and exhibits an architecture with side-by-side helical stacks, typical of functional RNAs such as riboswitches and ribozymes. The closed structure is stabilized by Mg^2+^ and by two tertiary contacts: tetraloop/tetraloop-receptor (TL/TLR) and metal-core/metal-core receptor (MC/MCR) (Figure 1B, Supplementary Figure 2A and Supplementary Text). The open state, by contrast, has no single structure; it consists of an ensemble of conformations in which the tertiary contacts are disrupted or are partially formed. Therefore, P4-P6 is a well-established model system for the study of RNA conformational ensembles, possessing an architecture that is broadly representative of functional RNAs and an ensemble that can be modulated with Mg^2+^ and mutations at defined tertiary contacts.

**Figure 1.**
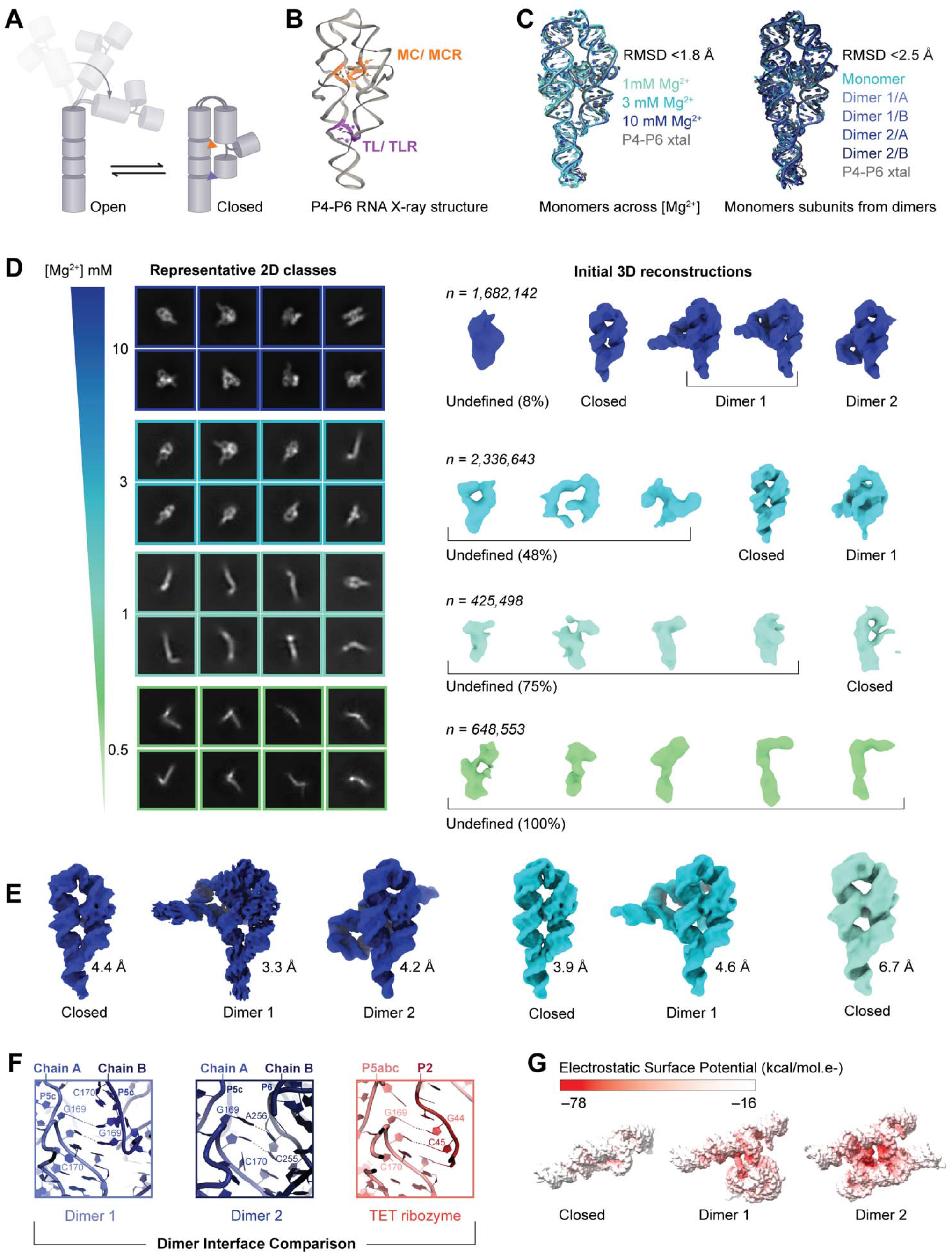
Stable P4-P6 RNA conformations revealed by cryo-EM. **A.** The P4-P6 domain of the *Tetrahymena thermophila* group I intron is a well-established model system that provides generalizable insights into RNA folding. The conformational landscape of P4-P6 is primarily defined as the interconversion between a compact “closed” state and an extended “open” state. **B.** The crystal structure (PDB ID: 1GID), shown as a cartoon representation, highlights the metal core–metal core receptor (MC/MCR) motif (orange) and the tetraloop–tetraloop receptor (TL/TLR) interaction (purple). **C.** RMSD comparisons of P4-P6 3D closed fold across [Mg^2+^] (left) and dimeric states (right), showing that the overall P4-P6 monomer fold is largely conserved. **D.** Cryo-EM data processing, outlining representative 2D classes and initial 3D reconstruction volumes at 10, 3, 1 and 0.5 mM Mg^2+^. **E.** Final 3D reconstructions of the two dimeric states and the closed monomeric state of P4-P6 RNA following iterative classification and refinement at varying Mg²⁺ (color coded as in D). **F.** Dimer 1 forms via a symmetric G169-C170 kissing-loop in P5c, while dimer 2 forms via non-canonical G169–C170 contacts with the partner subunit’s P6 helix, generating two equivalent interfaces. These residues also mediate long-range tertiary contacts with the P2 stem in the full-length *Tetrahymena* ribozyme (PDB: 7EZ0). **G.** Electrostatic potential mapping of the closed monomer and dimers.

### 3D reconstructions resolve stable states but fail to capture flexible states most populated at physiological [Mg^2+^]

We first used single-particle cryo-EM to reconstruct the stable states of P4-P6 across different [Mg^2+^] (Figure 1D). We collected data of P4-P6 folded at 10, 3, 1 and 0.5 mM Mg^2+^, in a constant background of 100 mM K^+^ (Supplementary Text). Despite the small size and dynamic nature of P4-P6, we determined multiple 3D reconstructions of distinct states at resolutions ranging from 3.3 to 6.7 Å (Figure 1D; Supplementary Figures 3-5).

At 10 mM Mg²⁺, *ab initio* 3D reconstruction yielded maps of the closed monomer and two distinct dimers (Figure 1D, dark blue). The closed monomer was refined to 4.4 Å (Figure 1E, dark blue) and matched the X-ray structure (RMSD ∼1.5 Å; Figure 1C, left). The two dimer states—hereafter referred to as dimer 1 and dimer 2—differ in their stabilizing interactions and in the relative orientation of the monomers (dark blue Figure 1D, E; Supplementary Figure 3). Dimer 1 was refined to 3.3 Å (Figure 1E, Supplementary Figure 3) and is stabilized by an intermolecular kissing loop, in which G169 and C170 in P5c of monomer A form reciprocal Watson-Crick base pairs with C170 and G169 of monomer B, orienting the two subunits at ∼76° (Figure 1E, Supplementary Figure 3E, 7B). Dimer 2 was refined to 4.2 Å. Like dimer 1, the interface is stabilized by intermolecular interactions involving G169 and C170 in P5c, forming non-canonical base pairs with C255 and A256 in the P6 stem of monomer B. Consequently, when the two dimers are superimposed on monomer A, monomer B in dimer 2 is flipped relative to its position in dimer 1, and the two subunits are oriented at ∼55° (Figure 1E, Supplementary Figure 3G, 7C).

The intermolecular contacts observed in dimer 2 correspond to those reported in the X-ray structure (*38*), whereas dimer 1 represents a configuration not previously described. In both dimers, their monomers adopt a fold superimposable with that of the isolated monomer (RMSD <2.5 Å, Figure 1C, right, Supplementary Figure 9), indicating that the fold is preserved across oligomeric states and consistent with the hierarchical nature of RNA. Illustrating the modularity of RNA, we discovered that G169 and C170 in P5c—the key residues forming intermolecular contacts in both dimers—are repurposing the exact tertiary binding interface used to engage the P2 stem in the full-length *Tetrahymena* ribozyme (Figure 1F) (*47*). The reuse of this interface, both in the context of the full ribozyme and the dimers, highlights the intrinsic propensity of this surface to form long-range interactions.

At 3 mM Mg^2+^, *ab initio* reconstruction yielded structures of the closed monomer and dimer 1 (Figure 1D, cyan). The closed monomer was refined to 3.9 Å (Figure 1E, cyan) with its atomic structure superimposable with that observed at 10 mM Mg^2+^ (RMSD ∼ 0.8 Å; Figure 1C). Dimer 1 was refined to 4.6 Å (Figure 1E, cyan; Supplementary Figure 4E) and its structure was similar to that at 10 mM Mg^2+^ (RMSD ∼0.96 Å; Supplementary Figure 9). The absence of a dimer 2 under these conditions suggests that Mg²⁺⁺ stabilizes the two dimers differentially, with dimer 2 requiring higher Mg²⁺ to reach a population sufficient for 3D reconstruction. Consistent with this, dimer 1 and dimer 2 differ in their calculated electrostatic potential, providing a structural basis for their distinct Mg²⁺ dependencies. Specifically, dimer 2 is more compact than dimer 1, having less solvent accessible surface area (Supplementary Figure 8) and buries a larger interfacial surface upon assembly (560 Å^2^, compared to 175 Å^2^ for dimer 1), thereby requiring stronger electrostatic screening to facilitate the tighter packing of dimer 2 (Figure 1G).

At lower Mg^2+^, obtaining well-defined 3D reconstructions was challenging as a larger fraction of particles occupied flexible conformations that cannot be aligned to a common reference map. The majority of 2D classes at 1 and 0.5 mM Mg^2+^ showed elongated, open conformations (Figure 1D; Supplementary Figure 5, 6). At 1 mM Mg^2+^, the closed state refined only to 6.7 Å (Figure 1E, Supplementary Figure 5C). At 0.5 mM Mg^2+^, the particles failed to produce interpretable 3D reconstructions (Figure 1C, light green), consistent with a broad, continuous distribution of open conformations that cannot be classified into discrete states.

The reconstructions above capture the discrete states that P4-P6 populates at each [Mg^2+^], but a substantial fraction of particles—exceeding 80% at low [Mg^2+^] (Figure 1D)—failed to converge to interpretable reconstructions. In standard 3D classification pipelines, such unaligned particles are routinely discarded, which often necessitates supra-physiological [Mg^2+^] to artificially restrict conformational mobility and force rigid state (*24, 48*). To overcome these limitations and quantify the full ensemble distributions, we turned to the raw 2D particle images, which reported on all conformations present in the distribution of picked particles, using the Bayesian reweighting approach described below.

### Ensemble reweighting infers compositional and conformational distributions from unclassified particle images

Our cryo-EM ensemble reweighting method (Figure 2) infers both compositional and conformational ensemble distributions of an RNA. The distribution is modeled as a nested mixture model (Methods), whose weights are directly inferred from raw, single-particle cryo-EM 2D projections, without particle alignment using simulation-based inference (*49*) and ensemble reweighting (*27*). It takes two inputs: a “prior ensemble” (Figure 2, bottom left) describing the allowed conformational and compositional space, and a set of unclassified cryo-EM particle images (Figure 2, top left), which we treat as independent and identically distributed (i.i.d.) samples from that space.

**Figure 2.**
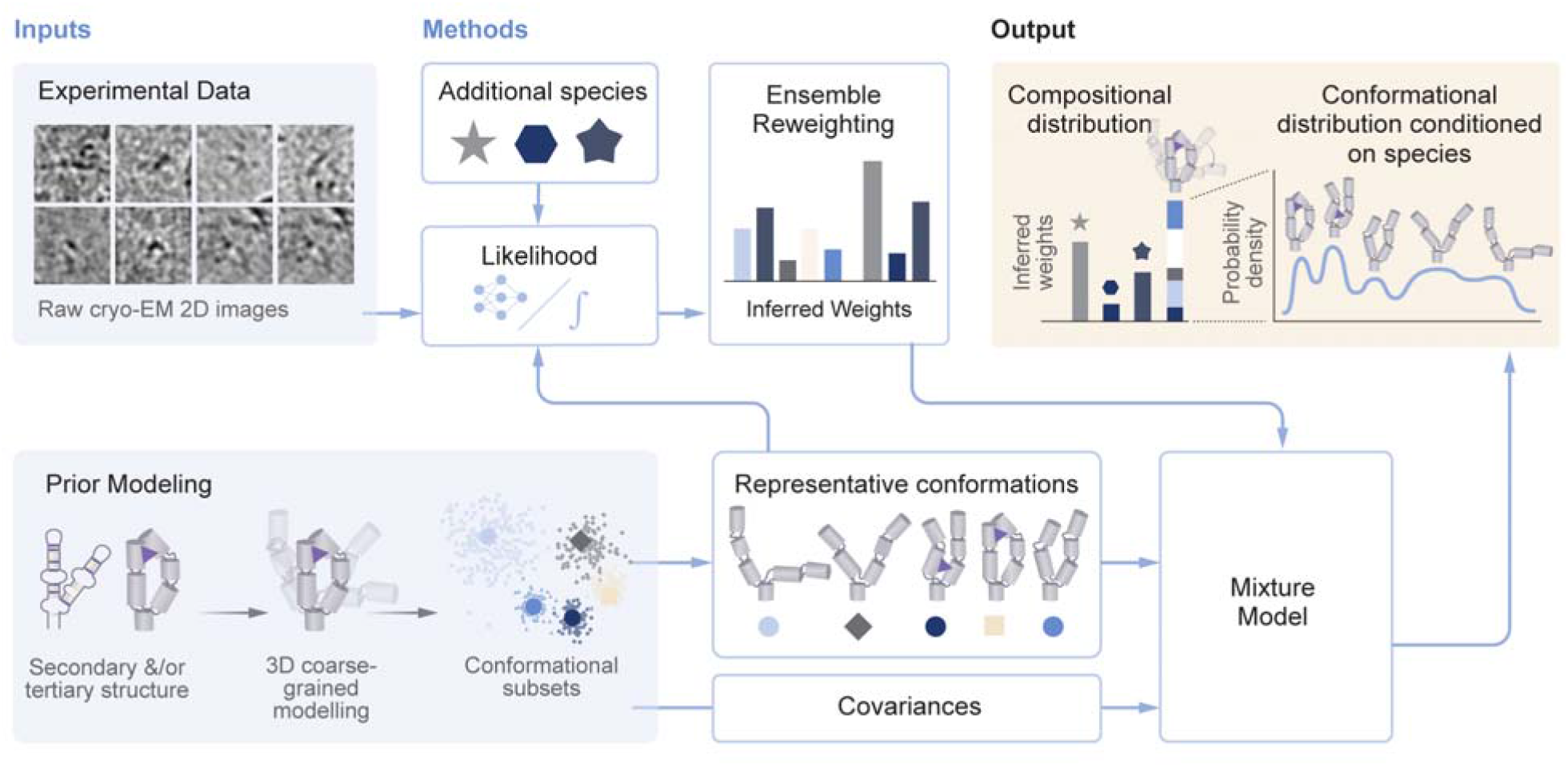
RNA cryo-EM ensemble reweighting. The method takes two inputs (blue boxes): (i) experimental cryo-EM particle images and (ii) a prior ensemble generated by coarse-grained sampling. The prior ensemble is clustered into conformational subsets, each of which contributes a representative conformation and a covariance. Additional compositional species (e.g., dimers) can be included. Representative conformations are compared with the particle images using an image-to-structure likelihood with either explicit or implicit pose marginalization. The resulting likelihood is used to infer probability weights via ensemble reweighting. These weights, together with the representative conformations and covariances, define the distribution as a mixture model. The output (beige box) is therefore a discrete distribution over compositional species and, conditioned on a given species, a continuous Gaussian mixture over conformations.

#### Building the Prior Ensemble

To generate the prior ensemble, we start from previously solved structures and employ coarse-grained (CG) simulations to sample accessible conformations. We leverage hierarchical folding and structural modularity, assuming a fixed secondary structure composed of stable helices connected by flexible junctions (Supplementary Figure 1). The RNA’s 3D space is dictated by the flexibility of junctions and the formation of modular tertiary contacts.

To make the image-to-structure comparison tractable and mitigate overfitting (*50*), we cluster similar conformations into subclasses (Supplementary Figure 10). From each subclass we extract a representative conformation, to which a probability weight is assigned during cryo-EM ensemble reweighting. The within-subclass covariance represents the flexibility around that representative and is used for calculating a Gaussian mixture model (GMM) that describes a continuous probability distribution (details below and in the Methods). For P4-P6, where discrete dimeric states were observed, we ran short CG simulations seeded from each dimer reconstruction, contributing one subclass per dimer to the prior ensemble. In this way, each source is leveraged for its core strength: simulations supply plausible local physical geometry and flexibility, while cryo-EM images quantify relative state populations.

#### Likelihood Calculations

For each experimental image and each representative conformation, we calculate an image-to-structure likelihood – the probability of observing that experimental image given the conformation. To evaluate the likelihood, we simulate thousands to millions of 2D templates from each representative conformation across poses and imaging parameters. Two different methods are used for likelihood calculations, cryoLike (*51*) and cryoSBI (*49*). In cryoLike, each experimental image is cross-correlated with all templates generated from a given structure and integration across templates yields the image-to-structure likelihood. In contrast to the explicit calculation of cryoLike, we use a modified version of cryoSBI, which amortizes the computational cost by learning a surrogate model of the likelihood. The surrogate likelihood consists of two neural networks: an embedding that reduces the cryo-EM image dimensionality, and a classifier that learns the probabilistic mapping between images and representative conformations (Supplementary Figure 11-14 and Supplementary Text). Once trained on simulated 2D templates from all representative conformations, cryoSBI evaluates image-to-structure likelihoods for experimental images at low computational cost (Supplementary Figure 15). Since the neural networks need only simulated images labeled with the representative conformation, the surrogate likelihood can also accommodate multiparticle templates, which we use to model the background and to define a “junk / aggregate” subclass during reweighting (Supplementary Figure 16 and Supplementary Text).

#### Ensemble Reweighting

The image-to-structure likelihoods are used to infer the weights for each conformation by maximizing the posterior probability (*52*). Here, we build on previous work (*27*) to include compositional heterogeneity (Supplementary Text). Total weights were calculated for each compositional state (e.g., monomer, dimer, junk/aggregate), by integrating across all conformations for each of these states.

For compositional states with substantial conformational heterogeneity, we modeled the conformational distribution using a GMM. The Gaussian centers and widths are obtained from the prior subclasses (Supplementary Text), and the mixture weights are those inferred for each subclass representative from the cryo-EM images. Importantly, the GMM lets us draw 3D conformations to visualize the reweighted ensemble (e.g., Supplementary Figure 17), and to compute key collective variables (CVs) over which the distribution can be projected.

In summary, the reweighted ensembles represent the compositional and conformational distributions observed in the cryo-EM sample (Figure 2, top right), enabling quantitative comparison across solution conditions, and estimation of ensemble-averaged observables.

### Reweighted ensembles capture Mg^2+-^dependent monomer-dimer and open-closed transitions

As described above, conventional single-particle cryo-EM analyses yielded high-to-moderate resolution reconstructions for three states of P4-P6: closed monomer, dimer 1, and dimer 2. However, these conformations only represent a relatively small number within a broad ensemble of possible conformations. Critically, at physiological [Mg^2+^], most particles populate open conformations that elude alignment and reconstruction. We therefore employed cryo-EM ensemble reweighting to obtain quantitative compositional and conformational distributions of P4-P6 across varying [Mg^2+^].

To perform ensemble reweighting, we prepared a separate, controlled series of cryo-EM experiments distinct from the dataset used for 3D reconstruction. P4-P6 was refolded at 10, 5, 3, 1, 0.5 and 0 mM Mg^2+^, in a background of 100 mM K^+^, and grids were prepared for each Mg^2+^ condition. To minimize variability from sample preparation or imaging, all samples were vitrified in a single session and imaged during the same microscope session (Figure 3A). Motion correction, CTF estimation, and micrograph curation were done identically for all datasets. Two different methods were used to pick particles in CryoSPARC (*53*), blob-picker and template picker (Supplementary Figure 18), which gave nearly identical reweighting results; the small differences observed are consistent with blob-picker retaining more junk particles. Three rounds of 2D classification were done, aiming to remove unambiguous junk while preserving all particles irrespective of conformation (Supplementary Text).

**Figure 3.**
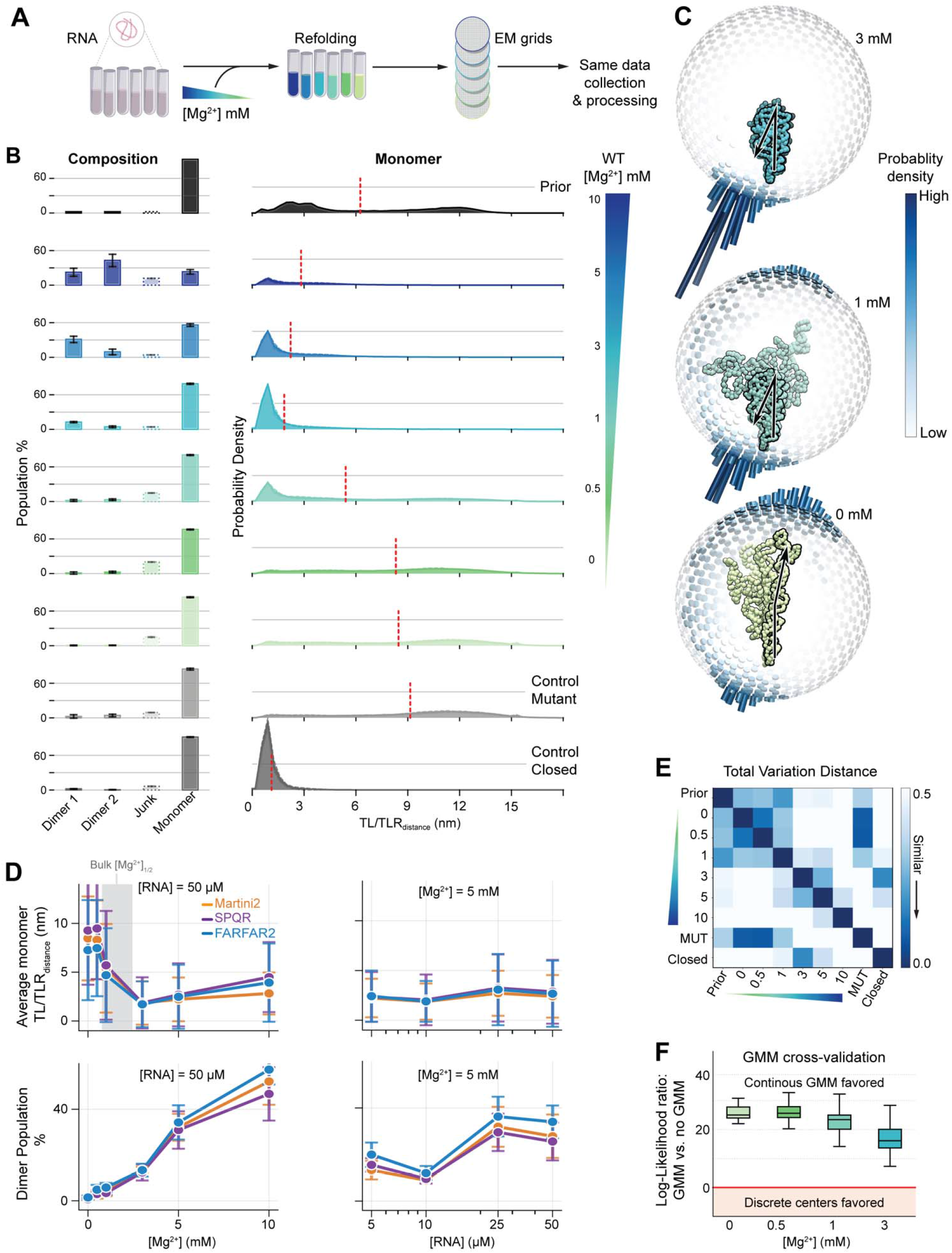
Quantification of Mg²⁺-dependent changes in the P4-P6 RNA ensembles using unclassified cryo-EM images. **A.** Schematic of sample preparation and data collection for ensemble reweighting. All samples are prepared, collected on the same days and processed with the same pipeline; the only variable is the [Mg²⁺] added to each tube. **B.** Ensemble reweighting results quantifying the Mg²⁺-dependent compositional distribution of P4-P6 RNA (left) and the continuous monomer conformational distribution along the TL/TLR_distance_ (right). Rows correspond to the prior, the wild-type titration series (10 to 0 mM Mg²⁺), and two controls. At the highest [Mg²⁺], dimers carry the highest weight; at 3 mM Mg²⁺ the closed state is highly occupied; at lower [Mg²⁺] a wide range of open conformations is populated. The controls are a mutant at 10 mM Mg²⁺ in which the tertiary contacts are disrupted, spanning wide-open conformations (MUT control), and a set dominated by closed conformations (Closed control). Red dashed lines mark average TL/TLR_distance_. **C.** Opening angular distributions of P4-P6 RNA at 3, 1 and 0 mM Mg²⁺, shown together with five representative conformations sampled from the GMM. Color indicates probability density. **D.** Averaged monomer TL/TLR_distance_ (top) and total dimer population (bottom) as a function of [Mg²⁺] at a constant [RNA] of 50 µM, for three prior ensembles (Martini2, SPQR, FARFAR2). The grey box in the top-left panel marks the bulk [Mg²⁺]_1/2_ range of 1-3 mM obtained from room-temperature experiments. The right-hand panels show the same observables as a function of [RNA] at a constant [Mg²⁺] of 5 mM: the average TL/TL_distance_ for the monomer distribution is insensitive to [RNA], whereas that of the dimer population is not. **E.** Similarity between probability distributions, measured by the total variation distance (TVD). Ensembles at adjacent low concentrations (0 and 0.5 mM) are mutually similar and similar to the mutant control, while 3 mM closely resembles the closed control. Darker colors indicate greater similarity (TVD = 0 for identical distributions). **F.** Cross-validation by the log-likelihood ratio between the continuous GMM and a distribution with the same weights placed on the discrete set of representative structures (|LLR|>3 is significant). At low [Mg²⁺] the continuous GMM outperforms the discrete representation; the ratio decreases with increasing [Mg²⁺].

The P4-P6 reconstructions were used as starting conformations to model the prior. To test the robustness of our results and its dependence on prior modeling, we sampled the 3D conformational space independently with three tools: Martini2 (*54*), SPQR (*55*), and FARFAR2 (*56*), generating tens of thousands of conformations that were then clustered for reweighting.

A single round of reweighting consisted of clustering the prior into forty monomer and two dimer subclasses, taking the cluster centers as representative conformations, and inferring their weights. To obtain statistics that report on robustness of landscape tessellation, we performed multiple rounds of reweighting: five rounds per Mg^2+^ condition, each round starting from a different clustering seed, so that the spread across rounds reported the statistical variability introduced by the discrete sampling of the prior (Supplementary Figure 19, 20 and Supplementary Text).

In comparison to 3D reconstructions, ensemble reweighting requires fewer input images to produce consistent results (*27*). Accordingly, twenty thousand randomly selected, unclassified cryo-EM particle images were used to reweight the representative conformations. This yielded a compositional distribution (monomers, dimer 1, dimer 2 and junk; Figure 3B, right) and a conformational probability distribution, represented by the GMM, across the Mg^2+^ titration series. The continuous GMM distribution was projected onto a CV, calculated as the distance between the TL and the TLR (TL/TLR_distance_; Figure 3B, left). All inferred distributions were consistent across different priors, clustering techniques, numbers of clusters, particle pickers, likelihood calculators and number of particles (Supplementary Figures 21-26).

As expected from the known strong role of ions on RNA folding, the inferred probability distributions were highly dependent on [Mg^2+^] (Figure 3B, top). At 10 mM Mg^2+^, dimers dominate over monomers (∼60% vs 25%, respectively). As we decreased [Mg^2+^], the population of dimers decreases, becoming undetectable at ≤1 mM Mg^2+^. Consistent with expectations, the population of dimers was a function of RNA concentration ([RNA]) while the monomer distribution was not (Figure 3D, left and Supplementary Figure 27). In agreement with dimer 2’s higher negative electrostatic potential (Figure 1F), dimer 2 is dominant over dimer 1 at 10 mM Mg^2+^, whereas dimer 1 was more populated relative to dimer 2 at 3 and 5 mM Mg^2+^. The cryoSBI junk (aggregate) class acted as a “rubbish collector” of high-order aggregates or ice contamination and absorbed approximately 10–30% of the particle weights (Supplementary Text, and Supplementary Figure 16), independently of [Mg^2+^].

At 3 mM Mg^2+^, the monomer conformational distribution showed a peak containing ∼70% of the monomer weight, centered at a TL/TLR_distance_ of ∼1 nm, corresponding to the closed state (Figure 3B, C). This distribution mirrored that of a control dataset, filtered beforehand to contain only particles that classify into the closed state (Figure 3B, E). At 1 mM Mg^2+^, the closed-state peak remained visible, but its total weight was substantially reduced (∼35%), and a substantial fraction of the weight is distributed across larger TL/TLR_distance_ values. As a result, the average TL/TLR_distance_ shifts from ∼2 nm at 3 mM Mg^2+^ to ∼6 nm at 1 mM Mg^2+^ (Figure 3D). At 0.5 and 0 mM Mg^2+^, no clear peak was observed around the closed state; instead, the weight was broadly distributed across TL/TLR_distance_ values, with an average of ∼9 nm. These distributions closely matched one obtained at 10 mM Mg^2+^ with a P4-P6 mutant (MUT) (Figure 3B, E), engineered to populate open conformations regardless of [Mg^2+^] conditions (Supplementary Figure 28) by removing the TL/TLR and MR/MCR contacts.

### The cryo-EM-derived [Mg^2+^]_1/2_ is consistent with solution measurements

The distributions above are consistent with well-established principles of RNA folding: (i) Mg^2+^ stabilizes compact over extended conformations; (ii) multimers are more probable at higher [Mg^2+^] and at higher [RNA]; and (iii) Mg^2+^ sensitivity is dependent on the electrostatic potential, e.g., dimer 2 *versus* dimer 1 (*38, 41, 43, 46, 57, 58*). Importantly, these distributions yield quantitative estimates for average observables that can be compared to other experimental techniques. One such observable is [Mg^2+^]_1/2_, defined as the [Mg^2+^] at which both open and closed states are equally populated. To estimate this, we defined the closed state as the collection of conformations with TL/TLR_distance_ ≤ 3 nm. Accordingly, the [Mg^2+^]_1/2_ estimated from our reweighted ensembles lies at ∼1 mM (Supplementary Figure 29). This value agrees well with solution measurements of P4-P6 folding. Single-molecule Förster resonance energy transfer FRET (smFRET) experiments under solution conditions matching ours (50 mM MOPS, pH 8.0, 100 mM KCl, 23 °C) yield a [Mg^2+^]_1/2_ of ∼1.4 mM, interpolated from the reported folding equilibrium constants at 1–5 mM Mg^2+^(*19*). At higher monovalent salt (200 mM NaCl, pH 7.0), smFRET and hydroxyl radical footprinting gave [Mg^2+^]_1/2_ values of 2.6 ± 0.2 and 2.4 ± 0.1 mM, respectively (*59*), whereas in the absence of monovalent salt, native gel electrophoresis gave 0.67 ± 0.03 mM (*45*). Our cryo-EM estimate therefore falls within the range of solution values and follows the expected trend of increasing [Mg^2+^]_1/2_ with increasing monovalent salt, which competes with Mg^2+^ for the RNA “ion atmosphere” (*58*). To our knowledge, this is the first equilibrium folding observable derived directly from cryo-EM particle images that can be quantitatively compared with solution measurements. This agreement supports the model that vitrified cryo-EM ensembles can report on Boltzmann-distributed solution ensembles, as recently supported by MD simulations (*29*) and discussed in more detail below.

### Continuous ensembles outperform discrete structures in describing flexible states

At low [Mg^2+^], the ensemble is dominated by high entropy states which are difficult to capture with a few discrete structures. To evaluate whether the prior-derived GMM covariances improve prediction of unseen images in these solution conditions, we performed cross-validation on a held-out set of three thousand images omitted from weight inference (Figure 3F). We quantified model performance using the log-likelihood ratio (LLR), comparing GMM samples (see examples in Supplementary Figure 17) against discrete, weighted cluster representatives, such as models sharing identical weights and centers, but lacking the covariance-derived flexibility around each center from the prior (Supplementary Text). Consistent with low [Mg^2+^] representing a high-entropy ensemble, the continuous GMM significantly outperformed the discrete representation at low [Mg^2+^] (LLR > 3, p < 0.05). In contrast, at higher [Mg^2+^], the LLR decreased as discrete cluster centers became sufficient to describe the more stable, well-defined conformations such as the closed state. The continuous GMM ensemble better predicts highly flexible open states on held-out images, extending analysis beyond discrete conformational models.

## DISCUSSION

RNA’s ability to fold into diverse 3D structures underlies its remarkable functional versatility in cells and viruses. However, our understanding of RNA structure-function relationships remains in its infancy, in part due to the dynamic nature of RNA structure which challenges conventional structural biology tools. By capturing individual molecules vitrified in near-native conditions, cryo-EM offers an unprecedented opportunity to dissect dynamic RNA ensembles (*12*). However, this potential has not yet been fully realized. Here, we combined single-particle cryo-EM reconstructions and a novel Bayesian reweighting framework to dissect the rich conformational ensembles of structured RNAs, using the paradigmatic P4-P6 RNA as a model system.

Cryo-EM reconstructions allowed us to visualize the stable monomer and multiple dimer states. Reconstructions across different [Mg^2+^] are qualitatively consistent with the well-established Mg^2+^-driven redistribution of states: more compact conformations concentrate negative charge and therefore, require higher Mg^2+^ to be sufficiently populated for 3D reconstruction. Generally, our analysis revealed a more complex conformational and compositional landscape than previously described for this model RNA. Despite these complexities, the reconstructions show that discrete structural domains behave as flexibly linked rigid modules (comprising stable secondary and tertiary structure motifs) whose architectures are conserved across Mg²⁺ conditions and oligomeric states. Our findings highlight how shared tertiary interfaces—such as the one mediating dimer 1—can be reused across distinct molecular states. Conventional cryo-EM reconstructions allowed us to visualize stable, well-defined states, but a substantial fraction of particles, particularly at physiological [Mg^2+^], occupy states that did not yield 3D reconstructions.

To move beyond the limitations of conventional cryo-EM, we developed a statistical framework for inferring RNA conformational distributions from unclassified cryo-EM images. Prior ensembles are constructed using the 3D reconstructions as starting states, and we leveraged the principles of hierarchical folding to computationally sample the allowed 3D conformational space: stable secondary-structure elements are connected by flexible junctions, whereas tertiary contacts are allowed to break. To account for compositional heterogeneity, the RNA ensemble was modeled as a nested mixture of compositional and conformational states.

Applied across datasets spanning different [Mg²⁺], the approach captures Mg²⁺-dependent conformational and compositional changes and recapitulates the bulk [Mg^2+^]_1/2_ transition for P4-P6 folding. At the lowest Mg²⁺, the inferred ensembles closely match that of a tertiary-contact deleted mutant measured at high Mg²⁺ and provide a better fit to held-out particles than a discrete set of conformations. Adding Mg²⁺ shifts the wild type toward compact states, and by 3 mM most particles occupy the closed monomer, mirroring the distribution obtained from a closed-state control dataset. At higher [Mg²⁺], dimers predominate. Consistent with a self-association equilibrium, the dimer populations depend on [RNA], whereas the monomer conformational distribution does not. These results are robust to parameter choices and prior force fields, enabling quantitative analysis of RNA ensembles.

Our statistical reweighting framework can provide uncertainty estimates, enabling quantitative comparisons between distributions, and cross-validation using held-out particle sets—capabilities that have largely been missing in cryo-EM heterogeneity analysis. In addition, the inferred ensemble defines a generative model from which we can draw structural samples. Looking ahead, we anticipate more expressive representations of ensemble probability densities, including neural network–based models, and ultimately the development of better forcefields and foundation models for RNA conformational ensembles that depend not only on sequence and structure, but also on environments such as solution conditions and temperature.

The cryo-EM RNA ensemble reweighting method relies on important assumptions. First, we construct the prior ensemble based on hierarchical RNA folding, assuming a fixed secondary structure. When alternative secondary structures might exist, however, tertiary folding can be modeled separately for each case, and a mixture model incorporating multiple secondary structures can be constructed. Second, we assume that the prior ensemble adequately spans the conformational and compositional variability present in the data. Verifying this assumption is challenging, and the development of systematic diagnostics remains an open problem. A complementary route would be to refine the prior iteratively against the experimental data (*60*), jointly optimizing the structures and their weights to extend coverage into regions of conformational space that the initial sampling missed. Third, we assume that the cryo-EM images are i.i.d. samples of the underlying ensemble. Upstream particle picking and preliminary classification can introduce selection biases, and the closed state control illustrates that heavily filtered particle sets produce inferred ensembles that mirror these biases. Moreover, cryo-EM datasets inevitably contain junk particles and aggregates, which can complicate inference. We found the inclusion of the explicit junk (aggregate) simulator in cryoSBI to be essential. This component acts as a statistical rubbish collector, absorbing weight from particles corresponding to aggregates or contamination that are intrinsic to cryo-EM data (Supplementary Text).

We show that ensemble properties of P4-P6 inferred from cryo-EM are consistent with bulk, room-temperature solution measurements. However, cryo-EM–derived ensembles may not coincide with room-temperature distributions for all systems, as the degree of correspondence depends on the freezing timescale relative to the lifetimes and relaxation dynamics of metastable states. Theoretical work suggests that when cooling is faster than relaxation, ensemble distributions remain similar to the equilibrium “hot” distribution. Further, even when relaxation is faster than cooling, recovery of the ensemble may still be possible by recording the measured ensemble at different cooling rates (*29*). In addition, grid preparation, the air–water interface, and day-to-day variability may influence ensembles measured by cryo-EM. For example, datasets collected on different days might show measurable discrepancies (Supplementary Figure 30). These considerations motivated us to adopt more controlled, biophysically guided data-collection protocols, in which all conditions are prepared and imaged on the same day for the ensemble reweighting (Figure 3A).

The achievable structural resolution of ensemble recovery remains an open question. Here, we focus on a system with large conformational changes that are distinguishable at low resolution; resolving subtler rearrangements will likely require more accurate matching of high-resolution features between templates and images and potentially more data. A key future direction is the development of improved neural-network representations of cryo-EM images, in which learned latent spaces enable more precise, high-resolution feature matching.

Functional RNAs are typically far more dynamic than the proteins most often studied by cryo-EM, and their dynamics are often central to their function. Capturing them requires extending beyond conventional 3D reconstruction, which resolves only relatively rigid states and discards particles that cannot be aligned. Our work advances single-particle cryo-EM from a tool for structure determination into a quantitative platform for mapping RNA conformational equilibria. By capturing flexible states that resist conventional alignment—the majority of particles under physiological conditions—this approach enables visualization of broad conformational landscapes, quantifies the relative populations of conformational and compositional states, and yields equilibrium observables that agree with orthogonal solution measurements. These capabilities provide a framework to resolve how sequence, architecture, and solution environment jointly govern the ensembles, and thereby the functions, of diverse RNAs.

## METHODS

### In vitro transcription, purification and folding of RNA

Templates for in vitro transcription were generated by PCR amplification of sequence-verified double-stranded DNA fragments (gBlocks; Integrated DNA Technologies). The sequences for all constructs and PCR primers are provided in Supplementary Table 4. The synthesized gBlocks have a ‘GG’ added at the 5’ end of the sequences of interest to facilitate transcription by T7 RNA polymerase. The transcription reactions comprised 8 mM of each NTP (ATP, UTP, CTP and GTP), 60 mM MgCl_2_, 30 mM Tris, pH 8.0, 10 mM DTT, 0.1% spermidine, 0.1% Triton X-100, 0.24 U/mL RNasin RNase inhibitor (Promega), and ∼0.14 mg/ml T7 RNA polymerase (prepared in house). The transcription reactions were incubated for 3 hours at 37 °C. The transcribed RNA constructs were ethanol precipitated, purified by denaturing gel electrophoresis (8% polyacrylamide), and buffer-exchanged into RNase-free water using 10 kDa cutoff Amicon centrifugal filters (Millipore). The RNAs were analyzed on a small denaturing gel for sample integrity. To fold the RNAs into their native conformation(s), the RNAs were resuspended at the desired concentration in a folding buffer solution containing 50 mM MOPS pH 8.0, 100 mM KCl and incubated at 50°C for 10 minutes. Approximately 0-10 mM MgCl_2_ was added to the samples, which were cooled to RT for 15 minutes. While sample preparation and data acquisition (see below) for the large datasets used for 3D reconstruction were conducted over multiple sessions, all datasets utilized for ensemble reweighting were prepared simultaneously and data was collected on the same microscope session (Figure 3A and Supplementary Text). This synchronization across all [Mg^2+^] was maintained to ensure consistency and minimize inter-dataset variability.

### Cryo-EM Grid Preparation and Data Collection Parameters

Cryo-EM grids were prepared using the standard procedure. Grids used were Quantifoil Carbon support Hole size - 1.2 mm, Spacing - 1.3 mm, Au mesh 300 (Electron Microscopy Services). Grids were cleaned on a Gatan Solarus Model 950 Advanced System, with these settings: Cleaning time - 7 seconds, Vacuum target - 70 mTorr, Vacuum Range - 0 mTorr, Pumping switch point - 20 Torr, Turbo Pump Speed - 750 Hz, Pumping timeout - 120 seconds, O_2_ Gas Flow - 8.7 sccm, H_2_ Gas Flow - 0.0 sccm, Ar Gas Flow - 26.3 sccm, Gas Flow Timeout - 20 seconds, Forward RF Target - 15 W, Forward RF range - 5 W, Maximum Reflected RF - 5 W, RF Tuning Timeout - 4 seconds, RF Tuning Attempts - 3. 3 µL of folded RNA solution (final RNA concentration - 50 µM, 50 mM MOPS, 100 mM KCl and 0-10 mM MgCl_2_) was applied on each grid and plunge frozen on FEI Vitrobot Mark IV (Thermo Fisher Scientific) using liquid ethane. The temperature of the Vitrobot Chamber was set at 10°C and humidity was set at 90 -100%. Blot time was set to 1 - 2 seconds, wait time to 0 seconds and blot force to 0. Filter paper was used for blotting (Ted Pella, Inc, Prod No. 47000-100). The grids were screened on the ThermoFisher Glacios Transmission Electron Microscope (TEM) operating at an accelerating voltage of 200 kV and equipped with a Falcon 4 direct electron detector (DED) at NYSBC. The best grids were chosen based on the particle distribution on ice and the number of optimal grid squares for imaging. Data collection was performed on ThermoFisher Krios TEM operating at an accelerating voltage of 300 kV and equipped with the Gatan K3 DED, BioQuantum GIF and/or ThermoFisher Scientific Cs corrector at NYSBC. Data were collected either at 0.334 Å/pixel (super-resolution mode) or at 0.844 Å/pixel (collection mode), with nominal magnification of 105,000X. Leginon (*61*) was used as data collection software. The list of Cryo-EM datasets for all conditions is provided in Supplementary Table 1-3. The movies contain 30-50 frames, collected at a dose of 55 - 56 electrons/Å^2^ and defocus range -0.5 to -2.5 µm.

### Processing of Cryo-EM datasets for 3D reconstruction

The Cryo-EM data were processed with CryoSPARC (*53*) using the Patch Motion Correction and Patch CTF Estimation tools. The micrographs were curated based on CTF, defocus, and ice contamination. For particle picking, a blob picker was initially run on a subset of 500 micrographs, followed by manual inspection. The particles were extracted using a box size of 384 pixels, and Fourier cropped to 128 pixels. Three rounds of 2D classification were performed, after which 2D class templates were used for template picking across the entire curated dataset. The extracted particles underwent 2 additional rounds of 2D classification. From there, an *ab initio* reconstruction was done with 5 classes, followed by 5 rounds of heterogeneous refinement to discard poor particles. The best particles were re-extracted at a full box size of 384 pixels, followed by one round each of homogeneous and non-uniform refinements. A tight mask was generated from the non-uniform refinement output and used in a final local refinement to obtain the final map. Resolution was estimated using the Gold Standard Fourier Shell Correlation at 0.143 cutoff (*62*). Details for cryo-EM data processing for each Mg^2+^ condition are detailed in the Supplementary Text, and Supplementary Figures 3-6. Data collection and map refinement statistics are reported in Supplementary Table 1.

### Structural Model Building

The P4-P6 crystal structure was docked into the monomeric and dimeric 3D maps using ChimeraX (*63*). The fitted models were then refined using molecular dynamics flexible fitting in Namdinator (*64*). Multiple rounds of real-space refinement were subsequently performed in Phenix (*65*) to resolve steric clashes and improve model geometry. MolProbity (*66*) was used for model validation. Model statistics are reported in Supplementary Table 1. RMSDs between P4-P6 monomers across Mg^2+^ and dimeric states were calculated in PyMOL relative to the P4-P6 crystal structure (PDB: 1GID) (Supplementary Figure 10). The interface angle between monomer subunits of the two P4-P6 dimers was calculated as the angle between the axes of the monomer subunits, in ChimeraX. The interfacial surface area of the two dimers was calculated using dr_sasa (*67*) (dr_sasa -m 1 -i <dimer pdb input> -chain A - chain B).

### The Ensemble Distribution

We adopted a nested mixture model to model the ensemble distribution. By treating the ensemble as a hierarchy, we first define discrete compositional species (e.g., oligomers or junk) via a categorical distribution. For the compositional species, we have the monomer, dimer 1, dimer 2 and an aggregate junk class simulated via cryoSBI (Supplementary Text). To capture broad and flexible conformational states, we model the ensemble distribution conditioned on the monomer as a continuous probability density using a GMM (Supplementary Text).

### Cryo-EM RNA Ensemble Reweighting

We employed Bayesian inference to reweight the prior ensemble to match the experimental cryo-EM distribution (*27*). By comparing individual, unclassified cryo-EM images against representative structures, the framework assigns a probability weight to each conformation. The original approach was extended to quantify discrete compositional heterogeneity and continuous conformational variability (Figure 2).

### Particles for cryo-EM Ensemble Reweighting

As noted above, all sample preparation was performed simultaneously, and data acquisition was conducted within a single microscope session for all Mg^2+^ concentrations utilized in the ensemble reweighting analysis. The Cryo-EM data were processed using CryoSPARC (*53*), with on-the-fly motion correction carried out using MotionCor2 (*68*) through the Appion (*69*) processing pipeline. The resulting motion-corrected micrographs were imported into CryoSPARC, where CTF estimation was performed using CTFFIND4 (*70*). Micrographs were curated based on a defocus range of -1 to -2.5 µm and a CTF fit between 3-12 Å, with astigmatism corrected to 0-0.3 µm for the wild-type P4-P6 dataset collected at 0 mM Mg^2+^. Particle picking was guided blob picking or by templates from representative structures derived from *CG* martini simulations of P4-P6 RNA and low pass filtered (Supplementary Text). This was applied to a randomly selected subset of 500 micrographs. Picked particles were manually inspected and initially extracted using a box size of 384 pixels and Fourier cropped to 128 pixels. Three rounds of 2D classification were performed to eliminate unambiguous junk particles. To improve particle centering, particles were re-extracted from the micrographs using a box size of 384 pixels, Fourier cropped to 128 pixels and restacked for processing.

### Prior Ensemble Modeling

To generate the prior ensemble, we tested different forcefields and modeling tools that capture the conformational variability from open to closed monomers as well as dimer flexibility. Since P4-P6 consists of several helices that form stable secondary structures, we model these as 8 rigid rods connected via flexible linkers that form dynamic tertiary contacts (Supplementary Figure 2A), enforcing these restraints for all force fields. First, CG molecular dynamics simulations were performed in GROMACS (*71*) with the Martini2 force field (*72*). Starting from the P4-P6 closed structure, we implemented an elastic network limited to secondary structure contacts, effectively creating eight rigid domains while allowing for tertiary flexibility. Monomer ensembles were generated by sampling starting conformations along a closed-to-open trajectory, followed by production runs totaling ∼100k conformations. For the dimer simulations, cryo-EM structures served as starting points, maintaining monomeric secondary structure restraints while leaving inter-monomer contacts unconstrained. See Supplementary Text for simulation details.

A complementary prior ensemble was generated using the SPQR software (*55*). This approach utilized eRMSD energy terms to enforce similarity to the reference structure across the same eight rigid domains (Supplementary Text), with conformational samples generated via Monte Carlo sampling, totaling ∼90k conformations. A similar constraining and sampling strategy was used to simulate the dimers starting from the cryo-EM structures.

Finally, we utilized fragment-based sampling via FARFAR2 (*56*) within the Rosetta suite (*73*). Initial sampling produced 100,000 structures of the P4-P6 monomer and the RNA strands near the 5’ and 3’ ends were constrained (with the -s flag) to ensure they formed the correct junction (residues 103-110, 211-217, 253-260). Even with this constraint, we found that the natively closed conformation was underrepresented and we supplemented the ensemble with samples constrained to the closed structure. Using the -rmsd flag, approximately 40,000 additional structures were generated across five distinct RMSD restraint tiers, ranging from 6 Å to 25 Å, to ensure a comprehensive sampling of the conformational landscape near the predicted closed state. The two cryo-EM dimer conformations were also sampled using FARFAR2. In addition to a 10 Å RMSD restraint relative to the cryo-EM structures, -obligate_base_pair flag was used across the monomer chains to enforce the tertiary contacts, giving rise to dimerization. For dimer 1, a cross-chain base pair was added between residue 169 in monomer 1 and residue 170 in monomer 2. For dimer 2, two cross-chain base pairs were added between residues 170 and 255 and residues 169 and 256. 100 structures of each dimer conformation were generated.

### Space Tessellation, Representative Structural Selection and Local Covariance Estimation

We tessellated the monomeric conformational space to extract structural representatives, and local covariance estimates from the prior ensembles. We primarily utilize Farthest Point Sampling (FPS) (*74*) to ensure expansive coverage of the CVs of interest, which include RMSD to the closed state and key tertiary contact distances (TL/TLR or MC/MCR), supplemented by K-means clustering to validate density-dependent clustering trends (Supplementary Text). The selected tile (or subclass) representatives serve as the fixed structural anchors used to compare with individual images via image-to-structure likelihood calculators (cryoLike or cryoSBI, see below and Supplementary Text). To obtain the local covariance estimates, we project all structures belonging to a tile into a reduced latent space via PCA. We ensure numerical stability and invertibility through Ledoit-Wolf shrinkage (*75*), which prevents rank-deficiency in sparsely sampled regions. As the dimers were well resolved by cryo-EM (Figure 1), each was treated as a single discrete subclass rather than a continuous GMM distribution.

### Image-to-Structure Likelihood Calculations

The ensemble reweighting hinges on the structure-image likelihood, which represents the probability that a specific structure is observed in the cryo-EM image. In this work, we calculate the likelihood using two distinct methods. To obtain a computationally efficient evaluation, we leverage cryoSBI (*49*) by training a surrogate likelihood on synthetic data, which offers the distinct advantage of amortization: once the model is trained, the structure-image likelihood for experimental images can be evaluated near-instantaneously without particle pose alignment. In contrast to the original cryoSBI framework, we train a classifier on a discrete set of structures to approximate the likelihood and employ a multi-particle simulator to account for background noise and a dedicated junk/aggregate class. Details about the simulation parameter ranges as well as the junk/aggregate simulator are presented in the Supplementary Text. We also utilize an explicit likelihood method, cryoLike (*51*), which computes the image-to-structure likelihood by integrating over poses using Fourier-Bessel functions (Supplementary Text).

### Weight Inference

We use the image-to-structure likelihood, from cryoLike or cryoSBI, as input to infer the ensemble weights via maximum *a posteriori* estimation using a multiplicative gradient algorithm (Supplementary Text) (*52*).

### GMM conformational samples

Conformations are drawn from the GMM and accepted only if they pass a bond-distance filter, which rejects sterically invalid proposals. This yields a continuous ensemble that retains local stereochemical validity (Supplementary Text and Supplementary Figure 17).

### Statistical Robustness and Error Estimation

To mitigate biases associated with the selection of specific representative structures, we employ a stochastic, multi-trial approach. The tessellation procedure is repeated across five independent iterations using different random seeds. This generates five distinct sets of ensembles (Supplementary Figure 19), allowing for the calculation of error bars that account for the uncertainty inherent in the discretization of the landscape. For each statistical reweighting round, one conformation per dimer was drawn at random from the prior simulations, restricted to those within 10 Å RMSD of the corresponding cryo-EM reconstruction.

### Ensemble Distribution Similarity

To quantify similarity between ensemble distributions, we used the Total Variation Distance (TVD). This metric measures the probabilistic difference between distributions on a scale from 0 (identical) to 1 (disjoint) (Supplementary Text).

### Cross-validation of GMM Ensemble on a Hold-out Set

To compare the out-of-sample predictive performance of the inferred ensembles, we employed a held-out-image cross-validation framework. For each experimental condition, a holdout set of 3,000 independent images was reserved to assess predictive performance via a LLR (Supplementary Text). By marginalizing over 50 structural samples from the GMM ensemble using the cryoLike likelihood function, we quantified how effectively it generalized to unseen data compared with the corresponding discrete ensemble with identical weights and representative centers. To ensure statistical robustness, we performed bootstrapping with replacement across both image and conformation axes over 50 iterations, providing a stable estimate of the LLR and confirming the GMM’s ability to capture heterogenous structural landscapes.

## DATA AVAILABILITY

Cryo-EM maps and structural models will be deposited in the Electron Microscopy Data Bank (EMDB) and Protein Data Bank (PDB) upon publication. Motion corrected micrographs and particles will be deposited in Electron Microscopy Public Image Archive (EMPIAR) and will become available upon publication. All the code used to process the data and generate results will be available upon publication.

## ACKNOWLEDGEMENTS

The Flatiron Institute is a division of the Simons Foundation. We acknowledge support from the CCB_X_ program of the Center for Computational Biology of the Flatiron Institute. This work was performed at the Simons Electron Microscopy Center at the New York Structural Biology Center, with major support from the Simons Foundation (SF349247). S.L.B. is a Howard Hughes Medical Institute (HHMI) Freeman Hrabowski Scholar and acknowledges support from the HHMI Hanna H. Gray fellowship, the Irma T. Hirschl/Monique Weill-Caulier Trust Research Award, and Stavros Niarchos Foundation (SNF) as part of its grant to the SNF Institute for Global Infectious Disease Research at The Rockefeller University. M.A is supported by a Gary Helman fellowship from The Rockefeller University. L.D. and R.C. acknowledge the support of the Goethe University Frankfurt and the Frankfurt Institute for Advanced Studies. This work was supported by the Deutsche Forschungsgemeinschaft (DFG, German Research Foundation) through the Cluster of Excellence SCALE – SubCellular Architecture of LifE (EXC 3094, Project ID (533751785) and the Collaborative Research Centre CRC 1507 (Project ID 450648163, project P09). We thank Lucy Reading-Ikkanda for assistance with figure preparation, and Sonya Hanson, Niko Grigorieff, Abhilash Sahoo and Simon Pobete for useful discussions.

