## Supplementary Information for "Quantifying RNA Folding Equilibria from Unclassified Cryo-EM Particle Images"

**4 Supplementary Figures** **25**

**5 Supplementary Tables** **52**

### 1 Supplementary Text: The P4-P6 RNA domain and cryo-EM experiments

#### 1.1 The P4-P6 domain

We use the 55-kilodalton P4–P6 domain of the *Tetrahymena thermophila* group I intron ribozyme, a well-characterized system that provides generalizable insights into RNA folding, electrostatics, and structural modularity. The P4–P6 RNA adopts a complex secondary structure (section 1.1.1), which can be approximated by an architecture consisting of two coaxially stacked helical bundles connected by a junction that induces an $\sim 180°$ turn, resulting in a parallel, globular fold typical of many riboswitches, ribozymes, and viral regulatory RNAs (Supplementary Figure 2A).

The P4–P6 RNA domain has served as a paradigm for RNA folding, characterized extensively by SAXS, X-ray crystallography, and single-molecule spectroscopy [1–7]. Its conformational landscape is defined by the transition between a high-entropy, heterogeneous open state and a compact closed state. This compaction is driven by two key long-range tertiary contacts: the tetraloop–tetraloop receptor (TL/TLR) and the metal core–metal core receptor (MC/MCR) interactions. While both monovalent and divalent cations promote the closed state through non-specific electrostatic screening, $Mg^{2+}$ ions specifically coordinate within the metal core to further stabilize the tertiary interface (Supplementary Figure 2A).

To modulate this equilibrium in the cryo-EM sample, we controlled the conformational transition using varying concentrations of $Mg^{2+}$ ions with and $K^{+}$ concentration constant at 100 mM. Additionally, we designed a mutant construct (MUT) to destabilize the closed state; this variant incorporates uridine-rich substitutions at the MC/MCR interface and a UUCG tetraloop mutation to disrupt the TL/TLR contact. We purified these P4-P6 variants using in vitro transcription and purified them by denaturing PAGE (detailed in Methods - Main Text), Supplementary Figure 2B. To visualize the resulting ensembles inferred from the cryo-EM images, we project the distributions onto collective variables (CVs) defined by the inter-residue distances of these two primary tertiary contacts.

##### 1.1.1 P4-P6 secondary structure

The P4-P6 secondary structure follows RNA base pairing rules and forms the P4, P5, and P6 helices connected by junctions, that position the helices for tertiary contact formation. Following the principles of RNA hierarchical folding, we define “helical blocks” in the P4–P6 RNA, shown as gray boxes (Supplementary Figure 2). In this context, helical blocks refer to modular A-form double helical segments that serve as rigid structural units within the RNA architecture. This representation reflects the hierarchical organization of RNA structure, where individual helices act as relatively rigid scaffolds connected by junctions, loops, and tertiary interactions. The connectivity between these elements introduces flexibility that enables the RNA to fold into well defined 3D shapes. The secondary structure of P4-P6 is shown in Listing 1.

**Listing 1** P4-P6 secondary structure in dot-bracket notation

```
....((((((...((((((.....(((.((((.(((..(((((((((....)))))))))..((.......))....)
    ))......)))))))....))))))..)).))))((...((((...(((((((((...)))))))))..))))
    ...))..
```

#### 1.2 Cryo-EM sample preparation and experiments

##### 1.2.1 Cryo-EM datasets for 3D reconstructions

To characterize the stable conformations of P4-P6 RNA, cryo-EM datasets were collected at four $Mg^{2+}$ concentrations (10, 3, 1, and 0.5 mM), with each dataset acquired on different days from independent sample preparations. Single-particle analysis was performed using CryoSPARC [8]. Supplementary Figures 3-6 present the cryo-EM processing and more details about the reconstructions steps for each dataset are presented in the next section.

##### 1.2.2 Cryo-EM 3D reconstructions

**Cryo-EM 3D Reconstruction of WT P4-P6 RNA at 10 mM $Mg^{2+}$.** A total of 7,880 movies were collected and processed in cryoSPARC [8] using Patch Motion Correction and Patch CTF Estimation. 7,730 micrographs were curated to exclude damaged grid regions or those with excessive ice contamination, retaining micrographs within a resolution range of 1.6–9.4 Å and a defocus range of -0.5 to -2.5 $\mu$m. Initially, blob picker was run on a randomly selected subset of 50 micrographs, followed by manual inspection of particle picks and extraction using a box size of 384 pixels, Fourier cropped to 128 pixels. After three rounds of 2D classification to remove poor-quality particles, the resulting 2D class averages were used as templates for template-based picking on the full curated dataset. Three additional rounds of 2D classification were performed to discard junk particles, resulting in $1,682,142$ particles. An *ab initio* reconstruction was performed with five classes, followed by five rounds of heterogeneous refinement to further remove junk particles and isolate the best particle sets representing the closed monomer, dimer 1, and dimer 2 separately. Particles representing each class were re-extracted with a box size of 384 pixels. One round each of homogeneous and non-uniform refinement was performed per class. To generate the final 3D maps, a tight mask was created from the non-uniform refinement results by optimally setting the map threshold and applying a dilation radius of 6 pixels and a soft padding radius of 6 pixels for local refinement. The final maps of the closed monomer, dimer 1, and dimer 2 were determined at resolutions of 4.4, 3.3, and 4.3 Å, respectively, as calculated using the gold-standard Fourier shell correlation (FSC) criterion at 0.143 (Supplementary Figure 3).

**Cryo-EM 3D Reconstruction of WT P4-P6 RNA at 3 mM $Mg^{2+}$**. A total of 19865 movies were collected and processed in cryoSPARC [8] using Patch Motion Correction and Patch CTF Estimation. 15,546 micrographs were curated to exclude damaged grid regions or those with excessive ice contamination, retaining micrographs within a resolution range of 2-10 Å and a defocus range of -0.5 to -2.5 $\mu$m. Initially, the blob picker was run on a randomly selected subset of 500 micrographs, followed by manual inspection of particle picks and extraction using a box size of 384 pixels, Fourier cropped to 128 pixels. After three rounds of 2D classification to remove poor-quality particles, the resulting 2D class averages were used as templates for template-based picking on the full curated dataset. Three additional rounds of 2D classification were performed to discard junk particles, resulting in $2,336,643$ particles. An *ab initio* reconstruction was performed with five classes, followed by five rounds of heterogeneous refinement to further remove junk particles and isolate the best particle sets representing the closed monomer and dimer 1 separately. Particles representing each class were re-extracted with a box size of 384 pixels. One round each of homogeneous and non-uniform refinement was performed per class. To generate the final 3D maps, a tight mask was created from the non-uniform refinement results by optimally setting the map threshold and applying a dilation radius of 6 pixels and a soft padding radius of 6 pixels for local refinement. The final maps of the closed monomer and dimer 1 were determined at resolutions of 3.9 and 4.6 Å, respectively, as calculated using the gold-standard FSC criterion at 0.143 (Supplementary Figure 4).

**Cryo-EM 3D Reconstruction of WT P4-P6 RNA at 1 mM $Mg^{2+}$.** A total of 4,054 movies were collected and motion corrected. CTF was estimated using CTTFFIND4 [9] in CryoSPARC [8]. 2,882 micrographs were curated to exclude damaged grid regions or those with excessive ice contamination, retaining micrographs within a resolution range of 3-20 Åand a defocus range of -0.5 to -2.5 $\mu$m. Initially, the blob picker was run on a randomly selected subset of 500 micrographs, followed by manual inspection of particle picks and extraction using a box size of 384 pixels, Fourier cropped to 128 pixels. After three rounds of 2D classification to remove poor-quality particles, the resulting 2D class averages were used as templates for template-based picking on the full curated dataset. Three additional rounds of 2D classification were performed to discard junk particles, resulting in $541,263$ particles. An *ab initio* reconstruction was performed with five classes, followed by five rounds of heterogeneous refinement to further remove junk particles and isolate the best particle sets representing the closed monomer and dimer 1 separately. Particles representing each class were re-extracted with a box size of 384 pixels. One round each of homogeneous and non-uniform refinement was performed per class. To generate the final 3D maps, a tight mask was created from the non-uniform refinement results by optimally setting the map threshold and applying a dilation radius of 6 pixels and a soft padding radius of 6 pixels for local refinement. The final maps of the closed monomer and dimer 1 were determined at resolutions of 3.9 and 4.6 Å, respectively, as calculated using the gold-standard FSC criterion at 0.143 (Supplementary Figure 5).

**Cryo-EM 3D Reconstruction of WT P4-P6 RNA at 0.5 mM $Mg^{2+}$.** A total of 3,986 movies were collected and processed in cryoSPARC using Patch Motion Correction and Patch CTF Estimation. 3,426 micrographs were curated to exclude damaged grid regions or those with excessive ice contamination, retaining micrographs within a resolution range of 2-10 Å and a defocus range of -0.5 to -2.5 $\mu$m. Initially, the blob picker was run on a randomly selected subset of 500 micrographs, followed by manual inspection of particle picks and extraction using a box size of 384 pixels, Fourier cropped to 128 pixels. After three rounds of 2D classification to remove poor-quality particles, the resulting 2D class averages were used as templates for template-based picking on the full curated dataset. Three additional rounds of 2D classification were performed to discard junk particles, resulting in $2,336,643$ particles. An *ab initio* reconstruction was performed with five classes, followed by one round of heterogeneous refinement with forced hard classification. Maps obtained were incomplete for further refinement, indicating that the particles are highly heterogenous which prevents proper particle alignment for 3D reconstructions (Supplementary figure 6).

##### 1.2.3 Structural Model Building

To examine the structural features of the P4-P6 maps, the P4-P6 crystal structure (PDB: 1GID) served as the starting model for fitting into both the monomeric and dimeric P4-P6 3D maps. The crystal structure PDB was docked into each map using ChimeraX, and the fitted models were subsequently refined using molecular dynamics flexible fitting (MDFF) via the Namdinator web server [10]. MDDF of the monomeric closed P4-P6 RNA was performed at 1, 3, and 10 mM $Mg^{2+}$ using map resolutions of 4.4, 3.9, and 6.7 Å, respectively, with additional parameters as follows: start and final temperatures of 298 K, G-force scaling factor of 0.2, 2,000 minimization steps, 20,000 simulation steps, 1 Phenix real-space refinement (RSR) cycle, and implicit solvent (GBIS) excluded. The resulting model was then imported into Phenix [11] alongside the cryo-EM map for additional real-space refinement, with secondary structure restraints applied throughout. Refinement was carried out using the 'minimization/-global', 'local/grid/search', and 'adp' strategies, with the following settings: maximum iterations of 50, 1 macro cycle, target bond RMSD of 0.0005 Å, and target angle RMSD of 1.5Å. Multiple rounds of Phenix real-space refinement were performed to resolve steric clashes and improve overall model geometry (Supplementary Figure 7A). The same workflow was applied to dimer 1 and dimer 2, with map resolutions adjusted to 3.3 and 4.2 Å, respectively (Supplementary Figure 7B and C). Model statistics for all structural models are reported in Supplementary Table 1.

##### 1.2.4 A control particle set for P4–P6 closed monomer

We used cryoSIEVE [12] to prepare a small high fidelity control dataset of closed monomer particles. CryoSIEVE is a post processing algorithm for single particle cryo-EM that iteratively removes particles contributing weakly to high resolution signal in the reconstructed map. Starting from a refined particle stack, the method performs repeated cycles of reconstruction, scoring, and particle removal, often retaining only a subset of particles while preserving or improving high resolution amplitudes and map resolution. CryoSIEVE was applied at the final stage of the cryoSPARC processing workflow to the larger P4–P6 RNA dataset collected at 3 mM $Mg^{2+}$, used for 3D reconstruction of the closed P4-P6 monomer. We first selected clean 2D classes corresponding to the closed P4-P6 monomer, yielding 40,122 particles. These particles were re-extracted with improved centering using a box size of 384 pixels and Fourier cropped to 128 pixels. One round of 2D classification, *ab initio* reconstruction, and homogeneous refinement was then performed to generate a low resolution mask. CryoSIEVE was subsequently run with parameters num_iter = 10, frequency_start =

40, frequency_end = 3, and retention_ratio = 0.8. This procedure yielded a reduced particle set enriched for the highest quality contributors to the reconstruction of the closed P4–P6 monomer. The final particle set contained 1,346 particles, reconstructed to 8.6Å, and serves as a control for subsequent ensemble reweighting analyses.

##### 1.2.5 Relatively small datasets are required for ensemble reweighting

While conventional cryo-EM workflows prioritize high-resolution structure determination at the expense of discarding large amounts of particles (e.g., see above), the comparative analysis of conformational ensembles across different datasets necessitates a more rigorous, non-filtering biophysical approach. Such methodology is essential to mitigate potential biases introduced during sample preparation and instrumentation, ensuring a faithful representation of the underlying molecular distributions for quantitative comparison.

An advantage of the cryo-EM ensemble reweighting is that fewer particles ($\sim 20,000$) are required to obtain statistical convergence of the ensemble distributions (Supplementary Figure 21). This can can be achieved by taking smaller cryo-EM datasets. Thus, we can interrogate RNA conformational landscape across multiple solution conditions in a single sample preparation and microscope session, offering direct biophysical views into how $Mg^{2+}$ modulates structural ensemble populations. Accordingly, cryo-EM grids were prepared on the same day (Main Text Figure 3A) using refolded WT RNA at $Mg^{2+}$ concentrations of 10, 5, 3, 1, 0.5, and 0 mM and mutant P4–P6 at 10 mM [$Mg^{2+}$], in a background of 100 mM KCl. To assess the influence of RNA concentration on structural ensemble populations, we additionally varied the concentration of WT P4–P6 RNA while maintaining $Mg^{2+}$ at a constant 5 mM. Datasets ($1,000$–$3,000$ micrographs) were collected per condition, in a single session on the same microscope (Supplementary Table 2-4). Micrographs were processed using on-the-fly motion correction with MotionCor2 [13] via the Appion processing pipeline [14]. The resulting motion-corrected micrographs were imported into CryoSPARC, where CTF estimation was performed using CTFFIND4 [9]. Micrographs were curated using selection criteria of defocus range $-1$ to $-2.5$ $\mu$m, CTF fit resolution $3 - 12$ Å and visual inspection for ice contamination. Particles were picked using either a blob picker or template picker (details below) on a randomly selected subset of 500 micrographs. After inspection, particles were extracted with a box size of 384 pixels and Fourier-cropped to 128 pixels. 3 rounds of 2D classification were performed to remove junk classes. The selected particles were then centered, re-extracted at a box size of 384 pixels, Fourier-cropped to 128 pixels, and restacked for subsequent likelihood estimation and ensemble reweighting.

**Particle picking:** Unbiased particle picking is a prerequisite for ensemble reweighting, where it is assumed that particles represent independent and identically distributed (i.i.d.) samples of the underlying distribution. For small, flexible RNA molecules like P4-P6, achieving this standard is particularly challenging, as low signal-to-noise ratios often cause background noise to be misassigned as true particles. While automated blob picking yielded a high density of approximately 1,650 picks per micrograph, this approach captured significant amounts of junk, aggregates or background particle picks. To improve picking efficiency, a template-based approach was adopted. 20 structures spanning the closed to open conformational states of P4-P6 RNA were selected from coarse-grained trajectories obtained using Martini2 (section 3.1.1). To minimize reference bias, molecular maps were low-pass filtered to 20 Å resolution using ChimeraX [15] prior to template generation in CryoSPARC, removing high-resolution features that could bias picking toward a specific conformation, while retaining sufficient low-resolution information to distinguish true particles from background noise. Particle picking using these templates yielded approximately 550 particles per micrograph, resulting in more selective recovery of P4–P6 particles over background, and representing an improvement in picking precision before NCC and power spectrum score filtering. Note that both picking strategies resulted in similar reweighting results with blob picker having slightly more weight for the junk and open state (Supplementary Figure 18).

### 2 Supplementary Text: The CryoEM ensemble distribution and ensemble reweighting

#### 2.1 The cryo-EM ensemble distribution

The ensemble distribution, $\rho$, specifies the probability that the RNA adopts a given *composition* and *conformation* under the sample's environmental conditions. In the limit of instantaneous vitrification, the cryo-EM ensemble can be related to the equilibrium free-energy landscape at the pre-freezing temperature via the Boltzmann distribution [16, 17].

To account for both compositional heterogeneity (discrete species) and conformational heterogeneity (continuous conformations), we represent the ensemble as a nested mixture model in which conformations are conditioned on composition. The compositional variable is modeled as

$$c \sim \text{Categorical}(\boldsymbol{\alpha}), \qquad \boldsymbol{\alpha} = (\alpha^{(1)}, \dots, \alpha^{C}), \qquad \sum_{c=1}^{C} \alpha^{c} = 1, \tag{1}$$

where $C$ is the number of compositional states and $\alpha^c$ denotes the population weight of the composition $c$, inferred from the cryo-EM images.

While discrete states are sufficient to model distinct compositional states, they are inadequate for capturing the behavior of entropy-driven, flexible open states. In these regimes, the system explores a continuous conformational landscape that cannot be faithfully represented by a sparse set of representative conformations. Therefore, for compositional states that exhibit substantial continuous conformational variability (e.g., the monomer), we model the conditional conformational density with a Gaussian mixture model (GMM - see details in section 2.3). For example, for the monomeric state ($c = 1$),

$$\rho(\mathbf{x} \mid \mathbf{c}=\mathbf{1}) = \sum_{\mathbf{m}=\mathbf{1}}^{\mathbf{M}^{(\mathbf{1})}} \alpha_{\mathbf{m}}^{(\mathbf{1})} \mathcal{N}\Big(\mathbf{x} \mid \boldsymbol{\mu}_{\mathbf{m}}^{(\mathbf{1})}, \boldsymbol{\Sigma}_{\mathbf{m}}^{(\mathbf{1})}\Big), \qquad \sum_{\mathbf{m}=\mathbf{1}}^{\mathbf{M}^{(\mathbf{1})}} \alpha_{\mathbf{m}}^{(\mathbf{1})} = \alpha^{(\mathbf{1})}, \tag{2}$$

where $\mathbf{x} \in \mathbb{R}^{\mathbf{3N}}$ denotes the Cartesian coordinates of a monomer conformation with $N$ atoms (or coarse-grained beads), $M^{(1)}$ is the total number of Gaussian mixture components in the GMM used to describe the monomer conformational landscape and $\alpha_m^{(1)}$ are mixture weights within the monomeric state, which add up to the total monomer population $\alpha^{(1)}$. The Gaussian means $\boldsymbol{\mu}_m^{(1)}$ correspond to representative prior conformations and $\boldsymbol{\Sigma}_m^{(1)}$ are the covariances that model conformational flexibility around each $\boldsymbol{\mu}_m^{(1)}$. In the following section, we describe a decoupled inference strategy in which the nested mixture weights are derived directly from the experimental data, whereas the GMM means and covariances are estimated from the prior monomeric ensemble (for details see section 3.1). We note that the continuous representation framework is not restricted to the monomer. It can be generalized to model conformational variability across diverse compositional states through an analogous formulation.

#### 2.2 Conformational and Compositional cryo-EM ensemble reweighting

The cryo-EM ensemble reweighting method [18] uses Bayesian inference and a prior ensemble to infer the ensemble distribution from individual cryo-EM images. Representative conformations are selected from the prior as described in section 3.1. Given a set of unclassified images (i.e., unbiased samples of the underlying distribution), the method infers the probability weight associated with each representative conformation, thereby reweighting the prior to match the ensemble supported by the data. Two requirements are essential: (i) the prior ensemble must cover the relevant compositional and conformational variability present in the images, and (ii) the images must be treated as unbiased samples of the cryo-EM ensemble, i.e., unclassified, indistinguishable (i.i.d. samples). We note that if particles used for the reweighting, were heavily classified and filtered using 2D or 3D classification cryo-EM methods, this can introduce selection bias and substantially distort the inferred ensemble weights.

In this work, we extend the original formulation [18] to jointly account for discrete compositional heterogeneity and continuous conformational heterogeneity within each composition. Consistent with our nested mixture model, we introduce a discrete compositional variable $c \in \{1, \ldots, C\}$ with $c \sim \text{Categorical}(\boldsymbol{\alpha})$, where $\boldsymbol{\alpha} = (\alpha^{(1)}, \ldots, \alpha^{(C)})$ and $\sum_{c=1}^{C} \alpha^{(c)} = 1$. For each composition $c$, we represent the conformational density using a set of representative conformations $\{\mathbf{x}_m^{(c)}\}_{m=1}^{M^{(c)}}$ (e.g., representative centers for the GMM), with associated nonnegative weights $\{\alpha_m^{(c)}\}_{m=1}^{M^{(c)}}$ satisfying

$$\sum_{m=1}^{M^{(c)}} \alpha_m^{(c)} = \alpha^{(c)} \qquad \text{for each } c \;, \tag{3}$$

as in Eq. 2 for the monomeric GMM.

Given a set of i.i.d. cryo-EM images $Y = \{y_i\}$, Bayesian inference yields the posterior over mixture weights

$$p(\boldsymbol{\alpha} \mid Y) \propto p(\boldsymbol{\alpha}) \prod_i \sum_{c=1}^{C} \sum_{m=1}^{M^{(c)}} \alpha_m^{(c)} \, p\Big(y_i \mid \mathbf{x}_m^{(c)}\Big), \tag{4}$$

where $p(y_i \mid \mathbf{x}_m^{(c)})$ is the image-to-structure likelihood and $p(\boldsymbol{\alpha})$ denotes a prior on the weights. We note that the mixture weights $\boldsymbol{\alpha}$ represent the relative occupancy of each compositional state within the experimental image set. Specifically, the probability that a given cryo-EM image $y_i$ originates from a particular compositional state.

For simplicity, in this work for the discrete compositional dimer classes, we set $M^{(c)} = 1$, i.e., we use a single representative conformation for each, as our 3D reconstruction analysis does not suggest significant conformational variability within each of these dimeric states (e.g., Supplementary Figure 3). Conversely, the monomeric state is modeled using a discrete set of conformations obtained by discretizing the prior ensemble, thereby capturing the range of relevant conformational changes (section 3.1.3). We evaluate $p(y_i \mid \mathbf{x}_m^{(c)})$ either by a surrogate likelihood using cryoSBI [19] that includes an additional junk class (section 3.2) or by explicit likelihood computation with cryoLike [20]. The inferred weights $\{\alpha_m^{(c)}\}$ are obtained by maximizing Eq. (4) (equivalently, a maximum a posteriori - MAP - estimate) using expectation–maximization (section 3.5).

#### 2.3 Modeling the continuous distribution of the monomer via a Gaussian Mixture Model

As introduced above, we approximate the continuous density conditioned on a composition using GMM defined in a coarse-grained Cartesian space. The continuous density for the monomer state ($c = 1$) is given in Equation (2) with $\boldsymbol{\mu}_m^{(1)} \in \mathbb{R}^{3N}$ mean coordinates of the $m$-th basin (or Gaussian centers), $\boldsymbol{\Sigma}_m^{(1)}$ the $3N \times 3N$ covariance matrix capturing the local structural fluctuations, and $\alpha_m^{(1)}$ the weight (population) of that specific conformational basin in the experimental ensemble.

Standard expectation-maximization approaches for fitting GMMs typically update both the weights ($\alpha_m^{(1)}$) and the spatial parameters ($\boldsymbol{\mu}_m^{(1)}, \boldsymbol{\Sigma}_m^{(1)}$) iteratively to maximize the likelihood of the observed data. However, optimizing the covariance matrices directly against the cryo-EM images presents severe computational and statistical challenges. Marginalizing the nuisance parameters at every iteration is computationally prohibitive, and estimating a full $3N \times 3N$ covariance from low-SNR projections is statistically underdetermined, leading our models to fit noise as apparent conformational variability.

Rather than estimating everything from the data, we decouple the two problems. MD force fields reproduce local geometry and fluctuations far better than they reproduce the relative populations of metastable states, which are sensitive to both the force field conditions. We therefore take the covariances from simulation and the mixture weights from the images: the simulations define where the wells are and how wide, the images define how much weight they have. Under the assumption of hierarchical folding, secondary structure elements remain intact and conformational variation is confined to junction flexibility. The priors therefore sample local fluctuations around each representative, capturing the extent of the landscape around it. We derive the local spatial parameters ($\boldsymbol{\mu}_m^{(1)}$ and $\boldsymbol{\Sigma}_m^{(1)}$) directly from the prior ensemble. Representative structural centers $\boldsymbol{\mu}_m^{(1)}$ are identified through a space tessellation, or clustering, of the prior ensemble and the local covariance matrices are initialized from cluster-specific variability of the prior and stabilized via a regularized approach that ensures numerical invertibility and avoids the computational overhead (see details in section 3.1.3). Then, we optimize the weights exclusively against the raw cryo-EM single-particle images (Section 2.2), ensuring that the recovered populations reflect the experimental ensemble distribution. Therefore, the experimental images dictate the underlying populations (the relative depths of the energy wells) while the physics-based simulations define the valid spatial geometries and structural variance within those wells. Crucially, by comparing the goodness of fit to unseen data (Section 3.8), we find that the GMM outperforms ensembles comprising just a discrete set of structures. This is particularly evident for experimental datasets with low $Mg^{2+}$ concentrations that exhibit highly heterogeneous states, as shown in Main Text Figure 3F.

#### 2.4 Representing and sampling the ensemble distribution

As described in Section 2.2, the inferred ensemble distribution is modeled as a nested mixture that accounts for both compositional and conformational heterogeneity. The compositional distribution is summarized by the inferred weights $\alpha^{(c)}$ for each species $c$ (monomer, dimer 1, dimer 2, junk), while the conformational distribution within each species is represented as the Gaussian mixture model of Eq. 2. Compositional probabilities, together with the across-set standard deviations obtained from the five independent ensemble sets defined in Section 3.1.2, are reported as bar plots in Main Text Figure 3B (left).

To visualize the high-dimensional conformational distribution of the monomer, we generate stochastic samples from the optimized GMM $\rho(x \mid c = 1)$ following the per-cluster sampling protocol of Section 3.1.3: component $m$ is selected with probability proportional to the inferred weight $\alpha_m^{(1)}$, a draw is made from its latent Gaussian, mapped back to Cartesian space through the inverse projection $W_m$, and accepted only if its backbone bond lengths are physically plausible. To preserve the relative monomer occupancy across experimental conditions, the number of accepted samples per condition is scaled by the inferred monomer weight, $N_{\text{samp}}(c = 1) \approx \text{int}(N_0 \sum_{m=1}^{M^{(1)}} \alpha_m^{(1)})$, with $N_0$ the total number of samples for each set. Samples are aggregated across the five sets and we estimated error bars based on the variance between sets (see below).

From each accepted sampled conformation, represented by one bead per residue, we evaluated the CVs of interest, e.g., the TL/TLR$_{\text{distance}}$ (A50–C120 BB2 atoms), and project the joint sample distribution onto the selected CV to obtain the one-dimensional marginal histograms shown in Main Text Figure 3B (right). The histograms are scaled so that their cumulative density encodes the inferred monomer population while their bin shape captures the conformational distribution within that state, allowing both compositional weight and conformational variability to be read off the same panel.

To visualize the folding of P4-P6 in 3D, we projected the monomer conformations onto a spherical histogram (Figure 3C Main Text). Two vectors were used to track the relative orientation of two major helices in P4-P6. The first vector was drawn along the largest helix in P4-P6, from the residues forming a hairpin turn (residues 237-140) at the bottom of the helix to the last residues forming a base pair at the top of the helix (residues 115 and 206). The second vector was drawn from the center of mass of the residues forming closing base pairs of the second major helix (residues 126 and 196 and residues 182 and 137). To draw the spherical histogram, we projected 20, 000 samples from the GMM calculated for different solution condition onto CVs defined by these vectors. That is, the polar coordinate on the spherical histogram is simply the angle between the vectors formed by the two helices for any given GMM sample. The azimuthal angle for the spherical histogram was chosen by comparing the orientation of the second helix in a given GMM conformation to the same helix in a reference folded conformation after both orientation vectors are projected into the x-y plane.

### 3 Supplementary Text: Computational Methods

#### 3.1 Prior Modeling

In this section, we provide a general description of how we construct RNA prior ensembles. Our ensemble model accounts for both compositional heterogeneity and continuous conformational variability. To model conformational heterogeneity, we build on principles of hierarchical RNA folding, in which stable secondary-structure elements are restrained and connected to each other by flexible junctions. During modeling, tertiary and intermolecular contacts are allowed to form and break. Within this framework, the prior ensemble is represented as a nested mixture that combines discrete compositional species (including monomer, dimer 1, dimer 2, and a junk/aggregation component with cryoSBI) with continuous conformational fluctuations for the monomer.

##### 3.1.1 Monomer continuous conformational modeling

To generate prior conformational ensembles for monomeric P4–P6, we used complementary modeling approaches at different resolutions, including Martini2 [21], SPQR [22], and FARFAR2 [23] (see below). Overall, our results are robust to the choice of force field, provided the prior ensembles cover the relevant regions of conformational space — a requirement met by all three models tested here.

Following the principles of RNA hierarchical folding, secondary-structure elements treated as restrained units (schematized as rigid rods in Supplementary Figure 2), while junctions and tertiary contacts were allowed to rearrange. Starting from the closed monomeric X-ray structure, we performed molecular simulations or Monte Carlo sampling with multiple random seeds to improve coverage. Since the tertiary contacts were not enforced, the open state was sampled in the ensemble. Hence, these ensembles span the full open-to-closed conformational landscape of the monomer (Supplementary Figure 10) and define the monomeric prior used for ensemble reweighting.

**Martini2.** We built the Martini2 force field [21] parameterization for RNA starting from the closed P4-P6 structure, PDB id 1GID chain A, using stiff network of elastic constraints. We used the P4-P6 secondary structure (Listing 1) to eliminate all such constraints that did not involve nucleotides forming secondary structure contacts. For the nucleotides involved in secondary structure elements, we kept the rubber constraints with successive neighboring residues and with the nucleotide to which the base pairing was formed. This resulted in the exclusion of all tertiary contacts from being enforced as constraints, and to 8 effective rigid elements, shown as rods in Supplementary Figure 2. All other RNA martini forcefield parameters, bonds, angles and torsions were kept the same as in ref. [21]. Molecular dynamics simulations were ran using the GROMACS software [24]. The starting conformation was embedded in an dodecahedron box with distance between the solute and the box of 1.4 nm, having $\sim$ 13000 water, antifreeze water $\sim$ 1500 and 157 NA atoms. For each simulation, we first performed a minimization and equilibration with 0.005 and 0.0005 ps time step respectively, and thermostat at 300K for 10000000 steps for the equilibration. This was followed by a short production run at 300 K for 25ns, and then one at 500 K for 500 ns with 0.005 ps time step. To sample the conformational space efficiently, we started from the closed monomer state ran two simulations to obtain closed-to-open starting positions of the monomer. Then, we randomly selected 10 conformations along the opening trajectory. We ran two production runs of 0.5 micro-s starting from each selected conformation with random initial velocities. Conformations were saved every 10000 steps, and trajectories from the final production runs were used as the Martini monomer prior ensemble, having 100k conformations over all runs.

For the dimer simulations, we used the models of the dimer as resolved by cryo-EM as starting structures and enforced the same secondary structure restraints as for the monomer. Note that the inter-monomer contacts were not enforced. The same MD protocols were used as with the monomer. We ran 10 replicas starting from only these cryo-EM structures. Conformations with RMSD to the staring structure of $< 10\mathring{A}$ were selected, and five conformations from this set were randomly selected to define each dimer subclass.

**SPQR**. We used the SPQR software to create a P4-P6 prior ensemble [22]. We employed the eRMSD metric to enforce structural similarity to a reference. This was achieved by calculating the RMSD relative to eight rigid blocks, which were defined by the secondary structure (Listing 1) and illustrated in Supplementary Figure 2. We used the SPQR MC function to run Monte Carlo (MC) simulations starting from the closed conformation at SPQR CG temperature 8, obtaining closed-to-open MC trajectories. We then randomly chose 10 conformations along the opening trajectory. We ran ten production runs of 1 billion steps long starting from each selected conformation with random initial seeds and all other SPQR parameters at default settings. Conformations were saved every 100k steps. For the dimer simulations, we used the cryo-EM dimer conformations as reference starting structures and enforced the same secondary structure restraints as for the monomer. Note that the inter-monomer contacts were not enforced. We ran 10 replicas starting from only these cryo-EM structures. Conformations with RMSD to the staring structure $< 10$ Å were used to define each dimer state.

**FARFAR2**. We utilized fragment-based sampling via FARFAR2 [23] to model the P4-P6 RNA by first generating an initial ensemble of 100,000 monomer structures. To ensure correct junction formation, of the 5' and 3' strands, residues 103-110, 211-217, and 253-260 were constrained using the -s flag with reference to previously resolved crystal structures of P4-P6 (PDB ID: 1G1D). Since the natively folded state remained underrepresented, this initial ensemble was supplemented with approximately 40,000 additional structures generated across five distinct RMSD restraint tiers (ranging from 6 Å to 25 Å) using the -rmsd flag. This let us thoroughly sample the conformational landscape near the predicted folded state. Additionally, the two cryo-EM-derived dimer conformations were modeled by generating 100 structures apiece. During these dimer runs, we enforced a 10 Å RMSD restraint to the cryo-EM structures. The -obligate_base_pair flag was also necessary to secure the cross-chain tertiary contacts required for dimerization. Specifically, dimer 1 was restrained with a cross-chain Watson-Crick base pair between residues 169 and 170, while dimer 2 required a cross-chain Watson-Crick base pair between residues 170 and 255 and a cross-chain Watson-Crick-Sugar base pair between residues 169 and 256.

##### 3.1.2 Monomer space tessellation and representative conformations

After simulating the conformational landscape of the monomer, we selected representative conformations with which to calculate image-structure likelihoods. To ensure broad coverage of the conformational space and to mitigate density biases from the prior, we primarily employed Farthest Point Sampling (FPS) [25], with K-means clustering and additional schemes evaluated as controls (detailed below). These cluster centers serve as structural templates for our forward models when calculating the image-structure likelihoods using (cryoLike [20] or cryoSBI [19]). As discussed in Sections 2.2 and 3.5, these likelihoods are used to infer the global mixture weights.

Concretely, a *cluster* $m$ is the Voronoi cell associated with center $\boldsymbol{\mu}_m^{(1)}$ in the two-dimensional CV space introduced below: it consists of all prior MD frames whose CV coordinates lie closer to $\boldsymbol{\mu}_m^{(1)}$ than to any other center under the Euclidean distance in this space. Each conformation in the prior ensemble is therefore assigned to exactly one cluster by this nearest-center rule, yielding a per-cluster sample of size $N_m$ that is further used to estimate the local covariance $\boldsymbol{\Sigma}_m^{(1)}$ in Section 3.1.3. Geometrically, the cluster centers act as anchors that demarcate the regions of conformational space over which probability weight is distributed within the cryo-EM ensemble.

The tessellation landscape was constructed on a two-dimensional CV space. For the primary results reported in the main text, the two CVs are (i) the RMSD to the closed-monomer X-ray structure after rigid-body alignment and (ii) the inter-residue distance between G108 and A183, which reports on the MC/MCR tertiary contact. FPS was applied directly to this 2D space using `quickfps` [25], returning $M^{(1)}$ centers that are maximally spread across the populated CV region. To assess how the inferred ensemble depends on the number of representative conformations, we repeated the procedure with $M^{(1)} \in \{5, 10, 20, 40, 80, 160\}$ (Supplementary Figure 23), which shows qualitatively consistent distributions. We adopted $M^{(1)} = 40$ as the primary value reported in the main text. This choice is consistent with the optimal ensemble resolution for weight inference obtained by maximizing the mutual information [26]. As control schemes, we additionally performed (a) K-means clustering on the same 2D CV space; (b) FPS on an alternative 2D CV space defined by the MC distance (G108-A183) together with the TL/TLR distance (A153-C223); and (c) a representative-resampling scheme in which the FPS partition was kept fixed but the per-cluster representative $\boldsymbol{\mu}_m^{(1)}$ was redrawn uniformly from within each cluster. For every scheme, the procedure was repeated with five independent random seeds, producing five sets of representative conformations whose spread provides our estimate of the uncertainty introduced by the choice of representatives, used downstream for reweighting and estimating the uncertainty of the populations along a given CV.

##### 3.1.3 Local covariance estimation for the GMM

Estimating the high-dimensional local covariance matrices, $\boldsymbol{\Sigma}_m^{(1)}$, is challenging because the $3N \times 3N$ Cartesian space (spanned by $N$ beads per conformation) is inherently rank-deficient and numerically unstable when estimated from the $N_m$ frames assigned to cluster $m$. To bypass this difficulty of dimensionality, we first rigid-body align all frames to a common reference using the rigid helical bundle, with residues $< 119$ or $> 203$, to remove global rotation and translation. We then project each cluster's structures into a reduced $D_m$-dimensional latent space ($D_m \ll 3N$) via adaptive Principal Component Analysis (PCA). Specifically, we eigendecompose the cluster-conditional empirical covariance and retain the leading $D_m$ eigenvectors to form the columns of a projection matrix $\mathbf{W}_m \in \mathbb{R}^{3N \times D_m}$. The truncation dimension $D_m$ is determined by the more restrictive of two criteria: (i) the minimum components required to reach 95% cumulative variance, or (ii) components with eigenvalues exceeding $10^{-3}\,\mathrm{nm}^2$. The second criterion matters for very tight clusters, where the variance is dominated by finite-sampling noise at displacements well below any physically meaningful scale.

Centered samples are mapped via $\mathbf{z} = (\mathbf{x} - \bar{\mathbf{x}}_m)\mathbf{W}_m$. Because we apply only this orthogonal rotation without standard-deviation rescaling, the latent samples remain decorrelated with empirical variances equal to the physical eigenvalues $\lambda_{m,i}$ (in $\mathrm{nm}^2$). Finally, we estimate the $D_m \times D_m$ latent covariance matrix $\widetilde{\boldsymbol{\Sigma}}_m^{(1)}$ using the Ledoit–Wolf shrinkage estimator [27]. Rather than enforcing strict diagonality, which would discard finite-sample off-diagonal residuals, the Ledoit–Wolf estimator shrinks the empirical matrix toward a scaled-identity target $\bar{\lambda}_m \mathbf{I}$ (where $\bar{\lambda}_m = \mathrm{tr}(\widetilde{\boldsymbol{\Sigma}}_m^{(1)})/D_m$). This dampens sampling noise along weakly populated directions and mathematically guarantees invertibility.

The corresponding Cartesian covariance is implicitly defined through the inverse projection, $\boldsymbol{\Sigma}_m^{(1)} = W_m\, \widetilde{\Sigma}_m^{(1)}\, W_m^{\top}$, and is never instantiated explicitly in the ambient $3N$-dimensional space. The resulting rank-$D_m$ Gaussian is supported on the cluster's PCA subspace and anchored at the representative structure $\boldsymbol{\mu}_m^{(1)}$. For heavily undersampled clusters where $N_m$ falls below the number of free covariance parameters in the latent space ($N_m \lesssim D_m^2/2$), we adopt a degenerate fallback: $\widetilde{\Sigma}_m^{(1)}$ is replaced by a tightly localized isotropic covariance ($10^{-6}\,\mathrm{nm}^2$ per direction) anchored at the representative structure $\boldsymbol{\mu}_m^{(1)}$, effectively collapsing the Gaussian onto the template until further samples warrant a proper estimate.

Although the PCA-latent representation ensures numerical stability and a well-conditioned $\boldsymbol{\Sigma}_m^{(1)}$, sampling directly from the Gaussian inevitably produces a fraction of proposals with unphysical local geometries (e.g., broken backbone bonds). To safeguard the physical validity of the final ensemble, we apply a generative bond-distance filter at the sampling stage. Each proposal is drawn from its latent Gaussian component, mapped back to Cartesian space through the inverse projection $x = z\,W_m^\top + \bar{x}_m$. Then, we measure the backbone bond lengths $\{d_i\}_{i=1}^{N-1}$ and score these against a Gaussian whose bond means $\boldsymbol{\mu}_{\mathrm{bond},i}$ and standard deviations $\boldsymbol{\Sigma}_{\mathrm{bond},i}$ are estimated once, from the full prior MD trajectory. Proposals whose mean bonds $\frac{1}{N-1}\sum_{i=1}^{N-1}|d_i - \boldsymbol{\mu}_{\mathrm{bond},i}|/\boldsymbol{\Sigma}_{\mathrm{bond},i}$ exceeds a cutoff of 4.0 are rejected as containing severe steric clashes or broken bonds. We oversample each cluster by a factor of 20 to maintain the target number of accepted samples after filtering.

With these procedures, we have the Gaussian centers given by the representative structures $\boldsymbol{\mu}_m^{(1)}$, estimates of the covariance for each subclass $\boldsymbol{\Sigma}_m^{(1)}$ conditioned on the monomeric species and conformational samples where some physical plausibility is ensured.

##### 3.1.4 Compositional species

Building upon the traditional cryo-EM pipeline in which distinct compositional states were identified, we modelled compositional heterogeneity as a discrete set of molecular species. For the dimeric species, we generated representative conformations using the same conformational modeling tools described above (Martini2, SPQR or FARFAR2) starting from the cryo-EM dimer structures and maintaining only restraints on secondary structure elements. Specifically, representative conformations for dimer 1 and dimer 2 were selected within 10 $\mathring{A}$ RMSD of their respective cryo-EM structures. This approach yielded conformations that capture fluctuations around each dimeric state, enabling uncertainty estimation for the inferred population weights. Consequently, each ensemble set consists of one representative randomly selected from the Dimer 1 prior, one from Dimer 2, $M$ monomeric representatives (selected using a clustering seed), and a junk/aggregation class modeled via the cryoSBI simulator. As noted above, five ensemble sets were generated using independent random seeds to account for uncertainty. Below, in section 3.2, we describe how we use the representative ensemble structures in cryoSBI to build an amortized classifier of the likelihood function.

#### 3.2 Image-to-structure likelihood calculation via cryoSBI

The central quantity required for the reweighting scheme described in section 3.5 is the structure-image likelihood $p(y_i \mid \mathbf{x}_m)$, the probability of observing image $y_i$ given that it contains a particle in conformation $\mathbf{x}_m$, marginalized over the unknown imaging parameters. Computing this quantity explicitly for hundreds of templates and hundreds of thousands of images is computationally intractable. Here, we describe how we can leverage cryoSBI [19] to approximate the structure-image likelihood using simulation-based inference (SBI). This allows us to amortize the evaluation of the likelihood, making the reweighing substantially more computationally efficient.

Let's start with one single particle image $y_i$ and a discrete set of candidate representative structures $\{\mathbf{x}_m\}$ with $m \in M = \sum_{c=1}^{C} M^{(c)}$ (note that here we include both conformational representatives and compositional species). For the image $y_i$ we assume that one of the structures $\mathbf{x}_m$ labeled by $m$ generated the image. CryoSBI allows us to learn an amortized surrogate model of the structure-image likelihood normalized over the candidate structures $\bar{p}(y_i|\mathbf{x}_m)$,

$$\bar{p}(y_i|\mathbf{x}_m) = \frac{p(y_i|\mathbf{x}_m)}{\sum_{m'=1}^{M} p(y_i|\mathbf{x}_{m'})} \ , \qquad p(y_i|\mathbf{x}_m) = \int p(y_i|\mathbf{x}_m, \boldsymbol{\phi})p(\boldsymbol{\phi})d\boldsymbol{\phi} \ . \tag{5}$$

Here, $p(y_i|\mathbf{x}_m, \boldsymbol{\phi})$ is the full likelihood, the probability of observing $y_i$ given the structure $\mathbf{x}_m$ and the nuisance imaging parameters $\boldsymbol{\phi}$ (particle orientation and translation, CTF parameters, signal-to-noise ratio, etc.), defined by the cryo-EM forward model 3.2.1. $p(\boldsymbol{\phi})$ is the prior over all the nuisance parameters $\boldsymbol{\phi}$. The normalized likelihood $\bar{p}(y_i|\mathbf{x}_m)$ differs from the pose-marginalized likelihood $p(y_i|\mathbf{x}_m)$ only by a factor that depends on the image $y_i$ but not on the structure $\mathbf{x}_m$. This factor is irrelevant for the purpose of ensemble reweighting, since we only need likelihood ratios to update the ensemble weights. Therefore, using $\bar{p}(y_i|\mathbf{x}_m)$ in place of $p(y_i|\mathbf{x}_m)$ in Eq. 6 yields exactly the same weights.

We use the cryoSBI framework with a classifier to approximate Eq. 5 (Supplementary Figure 11). We first generate a dataset $\mathcal{D}$ of simulated images with known structure labels. For each sample $i$, we draw a structure label $\mathbf{x}_{m_i}$ and imaging parameters $\boldsymbol{\phi}_i$ from the prior distribution $p(\mathbf{x}_m, \boldsymbol{\phi}) = p(\mathbf{x}_m) \cdot p(\boldsymbol{\phi})$, where $p(\mathbf{x}_m)$ is the prior over candidate structures, which we assume to be uniform. We then generate a synthetic image $y_i \sim p(y_i \mid \mathbf{x}_{m_i}, \boldsymbol{\phi}_i)$ with the forward model. Repeating this process $N$ times yields the dataset $\mathcal{D} = \{(y_i, \mathbf{x}_{m_i})\}_{i=1}^{N}$ of images annotated with their ground-truth structure, i.e. $N$ samples from the joint distribution $p(y, \mathbf{x}_m)$. On this dataset we train a classifier $q_\psi(\mathbf{x}_m \mid y) = \exp f_m(y)/\sum_{m'} \exp f_{m'}(y)$, with logits $f_m(y)$ and parameters $\psi$, to approximate the normalized structure-image likelihood $\bar{p}(y \mid \mathbf{x}_m)$. The parameters $\psi$ are obtained by maximizing the mean log-probability of the true labels, $\frac{1}{N}\sum_{i=1}^{N} \log q_\psi(\mathbf{x}_{m_i} \mid y_i)$, i.e. by minimizing the cross-entropy loss. This objective is simultaneously the neural posterior estimation (NPE) objective for a categorical density estimator and the multiclass classification objective of neural ratio estimation (NRE)[28, 29]. Consequently, after training the logits approximate the log-likelihoods up to a constant $c(y)$ that depends only on the image, $f_m(y) = \log p(y \mid \mathbf{x}_m) + c(y)$, so that their differences are the log-likelihood ratios $\log[p(y \mid \mathbf{x}_m)/p(y \mid \mathbf{x}_{m'})]$ between candidate structures.

Throughout, we assume that the simulator accurately reproduces real data. We validate this assumption a posteriori by comparing simulated and experimental image embeddings[19, 30] (Supplementary Figure 14).

##### 3.2.1 CryoSBI image simulator

We simulate cryo-EM images using a forward model based on the weak phase approximation (WPA), as described in ref. [19]. The three-dimensional density of each biomolecule is represented as a sum of Gaussian functions centered on each coarse-grained bead, where the Gaussian width $\sigma$ controls the spatial extent of each bead. A range of $\sigma$ values is used to capture variability in the electron density. Under the WPA assumption, the 3D density is projected onto a 2D plane to generate a noiseless projection of the electron density. This projection is then convolved with a contrast transfer function (CTF), which is parametrized by the following parameters: defocus $d$, B-factor $b$, and amplitude (AMP). The image center is shifted during the simulation, and Gaussian noise is added with the specified SNR, producing the final synthetic cryo-EM image $y$ used for training.

To simulate cryo-EM images containing several RNA molecules, the forward model is extended by an additional parameter $n_{\text{bg}}$, the number of particles in the background. The parameter $n_{\text{bg}}$ drawn is draw uniformly from $[n_{\text{min}}, n_{\text{max}}]$. Conformations for the background particles are independent samples from the same prior over conformations, orientations, and bead widths as the foreground particle (central particle). Image formation is carried out on a padded canvas that is center-cropped after CTF convolution, so that background particles can lie partially outside the final field of view. Particles are placed sequentially: the foreground particle is placed near the image center, with a small random in-plane shift drawn from a uniform prior of center translations, and each background particle is then proposed at a uniformly random position on the padded canvas and accepted only if it does not overlap any previously placed particle. Each structure is approximated by an enclosing ellipsoid whose principal axes are defined by the eigenvectors of the bead-coordinate covariance, with each semi-axis set to the maximum absolute projection of any bead along the corresponding axis. Each ellipsoid is rotated according to its particle's orientation and projected onto the image plane, and overlap between candidate placements is then evaluated by applying the Alfano–Greer criterion [**?** ] to the resulting ellipses. Candidates that fail this test after 1000 placement attempts are discarded.

***Junk/aggregate class.***

To allow classifiers trained on the simulated images to flag non-informative experimental particles such as ice contamination, aggregates, and junk picks, the simulator can optionally augment the training distribution with a "junk" class (see examples in Supplementary Figure16). Junk images are generated with the same probability as each of the conformational classes in the prior. Rather than containing the usual one foreground and $n_{\text{bg}}$ background particles, a junk image contains $n$ particles in total, with $n$ drawn uniformly from $[n_{\text{junk}}^{\text{min}}, n_{\text{junk}}^{\text{max}}]$. All $n$ particles are sampled independently from the same prior over conformations, orientations, and bead Gaussian widths. One particle is routed through the foreground image-formation pipeline so that the CTF and SNR scaling are applied as usual, while the remaining $n-1$ are placed as background using the same sequential overlap test described above. For P4-P6, we used the average radius of gyration ($\sim$ 60 Å) [4] to determine the maximum number of molecules that can fit within a structural class (containing a foreground particle) or a junk class. Given the box edge size of $\sim 256$ Å, a maximum of 4 particles can fit within the box without overlapping in the open state. Therefore, we define the aggregation class as having $n_{\text{junk}}^{\min} = 4$ and $n_{\text{junk}}^{\max} = 10$, while for the structural class, $n_{\text{bg}} \in [0, 3)$ resulting in a maximum total of 4 particles per box.

##### 3.2.2 CryoSBI priors

The following procedure outlines how samples are generated from the prior distribution $p(\mathbf{x}_m, \phi) = p(\mathbf{x}_m)p(\phi)$ to construct the dataset $\mathcal{D} = \{(y_i, \mathbf{x}_{m_i})\}_{i=1}^{N}$ used for training the surrogate model. The molecular structure $\mathbf{x}_m$ is sampled uniformly from the discrete set of $M$ structural representatives $\{\mathbf{x}_m\}_{m=1}^{M}$ obtained in section 3.1.2. This uniform prior over the representatives is a deliberate modeling choice: it provides the mathematical link between the cryoSBI posterior and the surrogate likelihood used in the reweighting step, allowing the trained posterior to be directly reinterpreted as a likelihood function. Particle orientations $q$ are drawn uniformly from the rotation group SO(3) to ensure isotropic coverage of all viewing directions. The remaining imaging parameters are sampled independently from uniform distributions over the ranges spanned by the training set: the Gaussian width $\sigma \in [0.5,\, 3.0]$ Å (up to $6.0\AA$ for models with only one bead per residue), the in-plane shift $s \in [-15,\, 15]^2$ Å (drawn independently along each axis), the defocus $d \in [0.5,\, 3.5]\,\mu$m, and the B-factor $b \in [1.0,\, 100.0]\,\text{Å}^2$. The amplitude contrast is fixed at AMP $= 0.1$. To cover a broad range of noise conditions, the signal-to-noise ratio is sampled log-uniformly, $\log_{10} \text{SNR} \sim \mathcal{U}\big(\log_{10}(5 \times 10^{-4}),\, \log_{10}(0.5)\big)$. The full set of imaging parameters is denoted $\boldsymbol{\phi} = \{q,\, \sigma,\, s,\, d,\, b,\, \text{SNR}\}$.

##### 3.2.3 CryoSBI training

The image-structure posterior is discrete, as each image is assigned a probability to one of the candidate conformations or junk class. To model this discrete image-structure posterior, we use a neural network classifier that predicts the probability $p(\mathbf{x}_m|y_i)$ of each structure $\mathbf{x}_m$ given an input image $y_i$. The classifier is trained using the cross-entropy loss, which is equivalent to maximizing the probability of the correct labels. During training, each batch of images is simulated on the fly, ensuring each image is seen only once and removing the need for a separate validation set. Training is performed with a one-cycle learning rate scheduler to achieve efficient convergence. The embedding network is a RegNetY [31] backbone with a 128-dimensional output, and the classifier is implemented as a prototype-based classifier [32]. We selected this combination based on a hyperparameter scan over 80 training configurations (Figure 12), in which the RegNet embedding paired with the prototype-based classifier head consistently achieved the lowest final training loss. We trained for 300 epochs with a batch size of 128, consuming 200 on-the-fly simulation batches of size 512 per epoch ($\sim 3.1 \times 10^7$ images in total)(for the exact training times see Tab. 5). The one-cycle scheduler peaked at a maximum learning rate of $5 \times 10^{-4}$, with weight decay $10^{-3}$ and gradient-norm clipping at 5.0.

##### 3.2.4 CryoSBI evaluation

After training, the model can be applied to experimental cryo-EM images. To ensure compatibility between simulation and experiment, the pixel size and image dimensions are matched beforehand. Each experimental image is preprocessed by whitening and normalization to align its statistical properties with those of the simulated training data. The network outputs logits for each structural label, which correspond to the discrete posterior log probabilities. These outputs can be used directly to construct the structure–image likelihood matrix, $p(y_i|\mathbf{x}_m)$, required for Bayesian reweighting of conformational ensembles as described below. The inference throughput averaged over all models and all datasets is 0.7 s per 1000 images. In comparison to explicit likelihood methods, this approach is computationally highly efficient during inference; however, it requires a computational investment during the training phase (Supplementary Figure 15).

#### 3.3 Explicit image-to-structure likelihood evaluation with cryoLike

An explicit likelihood calculation was also evaluated using the CryoLike software package [20], which builds on the BioEM method [33]. It provides a framework for comparing conformations to experimental images by working with a Fourier-Bessel representation and explicitly integrating-out the unknown image poses. For this analysis, we used the same an image descriptor, with a grid size of $128 \times 128$ voxels and a voxel size of 2.007 Å, as in section 1.2.5. To balance computational speed and physical accuracy, calculations were performed with single-point precision using a resolution factor of 0.4 and 128 in-plane rotations. The forward modeling of the Marini conformations utilized a 3.0Å Gaussian width for all CG beads, while for the cross-validation of the GMM samples we used one bead per residue centered at the BB2 bead with Gaussian width of 6Å. The viewing distance was set to $1.0/4\pi$ to standardize the projection geometry.

#### 3.4 Choosing between cryoSBI and cryoLike

The choice between these two methods comes down to a trade-off between upfront training time and per-image computation. As described above, cryoLike calculates the image-to-structure likelihood by explicitly comparing each experimental image against a fine grid of projections of each structure. In contrast, cryoSBI amortizes pose inference during a computationally intensive initial training phase, leading to much faster evaluation later. The computing costs for these two methods are compared as a function of the number of image evaluations and for fixed number of models in Supplementary Figure 15B for the parameters described above, where we find that for $< 10000$ images cryoLike is more computationally efficient while for more images cryoSBI is faster.

In terms of accuracy, we compare the reweighting results on synthetic data for both cryoLike and cryoSBI in Supplementary Figure 15A. We observe that for images with high noise levels, cryoSBI reweighting achieves a better recovery of the ground truth distribution. While cryoLike's performance can be improved by increasing the resolution factor, doing so significantly escalates the computational cost (Supplementary Figure 15B).

A key advantage of cryoSBI is its ability to handle more complex particle classes—such as junk or aggregates—by leveraging more sophisticated simulators. On the other hand, cryoLike is highly advantageous when evaluating new conformations; because it requires no training, and samples from the GMM can be easily evaluated on a small test set of images. Consequently, we present our main results and systematic testing using cryoSBI, while utilizing cryoLike for cross-validation.

#### 3.5 Weight inference for cryo-EM ensemble reweighting

As described above, cryoSBI or cryoLike were employed to estimate the image-to-structure likelihood, evaluating each representative conformation across all compositional species ($\mathbf{x}_m^{(c)}$) against every experimental image ($y_i$). These likelihood values subsequently serve as the foundation for the ensemble reweighting.

Following the theoretical framework detailed in section 2.2, we employ the posterior probability defined in Eq. 4 to determine the optimal weights, $\boldsymbol{\alpha}$, for the nested mixture model that represents the cryo-EM ensemble distribution, which is statistically consistent with the observed images given the image-to-structure likelihood matrix. Utilizing it in Eq. 4, we determine the optimal global weights $\alpha_m^{(c)}$ with decoupled expectation maximization updates. This is known as the Multiplicative Gradient (MG) algorithm [34], which formalizes the ensemble reweighting principles established in ref. [35]. MG provides an iterative framework to identify the weight vector $\boldsymbol{\alpha}$ that maximizes the log-posterior of the image dataset $Y$. By adopting a flat prior over the weight space $p(\boldsymbol{\alpha})$, the maximum *a posteriori* estimate becomes equivalent to the maximum likelihood estimate derived from the rescaled log-likelihood

$$\mathcal{L}(\alpha^{(c)}) := \frac{1}{I}\sum_{i=1}^{I}\log\left(\sum_{c=1}^{C}\sum_{m=1}^{M^{(c)}}\alpha_m^{(c)}p\Big(y_i \mid \mathbf{x}_m^{(c)}\Big)\right), \tag{6}$$

where we have divided by the total number of images $I$ to scale the objective with the number of images.

The MG algorithm is computationally efficient and proceeds by initializing the weights uniformly, consistent with a non-informative flat prior. In each iteration, the current weight vector $\alpha_m^{(c)}$ is updated to $\tilde{\alpha}_m^{(c)}$ via a coordinate-wise multiplication with the gradient of the log-likelihood: $\tilde{\alpha}_m^{(c)} := \alpha_m^{(c)}\frac{\partial\mathcal{L}(\boldsymbol{\alpha})}{\partial\alpha_m^{(c)}}$. No normalization is needed for the simplex constraint, as $\frac{\partial\mathcal{L}(\boldsymbol{\alpha})}{\partial\alpha_m^{(c)}}$ is always positive, and $\sum_c\sum_m\alpha_m^{(c)}\frac{\partial\mathcal{L}(\boldsymbol{\alpha})}{\partial\alpha_m^{(c)}} = 1$ for any choice of $\boldsymbol{\alpha}$ in the simplex. Thus, this multiplicative approach uses the structure of $\mathcal{L}(\boldsymbol{\alpha})$ to ensure that the updates remain on the probability simplex, iteratively refining the ensemble occupancy to maximize the objective function. There is no step-size selection, and the only user choice is a stopping tolerance parameter.

For all experimental datasets, we adopt the stopping tolerance parameter according to convergence criteria established in ref. [35], which typically results in termination within approximately 10 iterations. While this algorithm has appeared in various contexts—notably as the Richardson-Lucy iteration [36–39]—its convergence is well-characterized under mild conditions on the likelihood distributions [34, 39]. By deriving the full gradient $\nabla_{\boldsymbol{\alpha}}\mathcal{L}(\boldsymbol{\alpha})$ of the log-posterior in Eq. (6), it can be demonstrated that the MG update is mathematically equivalent to the M-step update for mixture weights in the expectation-maximization algorithm [35, 40].

The resulting weights are combined with the representative cluster centers and covariances to construct an approximate ensemble distribution via the nested mixture model and GMM. This cryo-EM reweighted ensemble provides a flexible representation of the compositional and conformational distribution, enabling sampling of conformations, projection onto low-dimensional coordinates, quantitative comparison of ensembles, and computation of ensemble-averaged observables. Below we describe some postprocessing steps that we can do to study and compare the inferred distributions across varying magnesium concentrations.

##### 3.6 Statistical Ensemble Reweighting

To quantify the uncertainty introduced by the selection of specific representative structures, the five independent ensemble sets generated in Section 3.1.2—each consistently spanning the relevant CV and compositional space, but built from a different random seed and thus comprising a different concrete set of representatives—are reweighted independently following the procedure of Section 3.5. The spread across these five reweighting outcomes provides our estimate of the sensitivity of the inferred populations to the choice of representatives.

For the compositional weights $\alpha^{(c)}$, we report the mean and standard deviation across the five sets, with the standard deviation taken as the across-set statistical uncertainty (error bars in Main Text Figure 3B, left). For the conformational distribution of the monomer, samples drawn from the per-set GMMs (Section 3.1.3) are pooled in the common CV space, so that the resulting histograms reflect the combined uncertainty arising from the choice of representatives, the weight inference, and the per-cluster sampling (Main Text Figure 3B, right - shaded regions).

##### 3.7 Ensemble Distribution Similarity

To compare cryo-EM ensembles inferred under different experimental conditions on a common footing, each ensemble is summarized as a single discrete probability distribution that jointly encodes its compositional and (monomeric) conformational content. Specifically, for each condition we (i) bin the pooled monomer CV samples along a chosen CV into $K = 5$ equal-width bins on a fixed range; (ii) normalize the resulting histogram to sum to unity and rescale it by the inferred monomer weight $\alpha^{(1)}$, so that the monomer bins sum to $\alpha^{(1)}$; and (iii) concatenate with the compositional weights $\alpha^{(\mathrm{Dim}_1)}$, $\alpha^{(\mathrm{Dim}_2)}$, $\alpha^{(\mathrm{junk})}$. The resulting probability vector $P$ has length $K + 3$ and sums to one, defining a discrete distribution over the joint sample space $\Omega = \{\text{monomer-CV bin}_1, \ldots, \text{monomer-CV bin}_K,\ \mathrm{Dim}_1,\ \mathrm{Dim}_2,\ \text{junk}\}$.

The disparity between two such distributions $P$ and $Q$ over $\Omega$ is quantified by the Total Variation Distance (TVD) [41],

$$d_{\mathrm{TV}}(P, Q) \;=\; \tfrac{1}{2} \sum_{x \in \Omega} |P(x) - Q(x)| \;\in\; [0, 1], \tag{7}$$

which equals 0 when $P$ and $Q$ coincide and 1 when they have disjoint supports. Because $\Omega$ couples compositional and conformational information by construction, $d_{\mathrm{TV}}$ simultaneously penalizes mismatches in species occupancy and in the shape of the monomeric CV distribution. We evaluate Eq. (7) pairwise across all conditions, separately for each CV of interest, and summarize the resulting similarity matrix as a heat map (Main Text Figure 4D).

#### 3.8 Cross-validation on held-out images

To evaluate the fidelity of our inferred GMM ensembles, we would like to ensure that the mixture model captures the intrinsic landscape rather than overfitting to stochastic noise inherent in the cryo-EM training images. Because maximum-likelihood approaches can be prone to overfitting in low signal-to-noise regimes, we validate our inferred ensembles by evaluating their predictive performance on unseen images (henceforth test image set). Therefore, for each experimental condition, we reserve a set of independent holdout test images, $Y_{\mathrm{test}} = \{y_i\}$.

We approximate the continuous GMM ensemble by generating samples $J$ from it, $X = \{\mathbf{x}_j\}_{j=1}^{J}$, where $x \sim \rho$. We calculate the likelihood of an individual image $y_i$ by marginalizing over the samples $\{\mathbf{x}_j\}$. To evaluate the predictive power of the model, we determine how well the samples explain these unseen images using the total log-likelihood over the test dataset,

$$\log p(Y_{\mathrm{test}} \mid X) = \sum_{i \in Y_{\mathrm{test}}} \log \frac{1}{J} \sum_{j} p(y_i|\mathbf{x}_j) \ , \tag{8}$$

where $p(y_i|\mathbf{x}_j)$ is the image-to-structure likelihood. Because the samples $\{\mathbf{x}_j\}$ do not necessarily correspond to the representative cluster centers used to train cryoSBI, we use the explicit likelihood calculation from cryoLike [18] to compute $p(y_i|\mathbf{x}_j)$.

To quantify the relative performance of one ensemble, $\mathbf{X}^a$ (e.g., GMM samples), against another, $\mathbf{X}^b$ (e.g., fixed structures), on a given test set $Y_{\mathrm{test}}$, we use the log-likelihood ratio (LLR) defined as

$$\mathrm{LLR}(Y_{\mathrm{test}}, \mathbf{X}^a, \mathbf{X}^b) = \log \left( \frac{p(Y_{\mathrm{test}} \mid \mathbf{X}^a)}{p(Y_{\mathrm{test}} \mid \mathbf{X}^b)} \right) \ . \tag{9}$$

A positive LLR ($> 0$) indicates that the ensemble $\mathbf{X}^a$ generalizes better than ensemble $\mathbf{X}^b$ to new unseen data and more accurately represents the true structural distribution of the sample, with statistical significance at $> 3$. If the LLR is negative then $\mathbf{X}^b$ better represents the unseen data.

The results presented in the main text aim to determine whether the GMM monomer ensemble, $\rho(\mathbf{x} \mid c = 1)$ defined Eq. 2, is a more effective representative of unseen data than an ensemble composed of a fixed set of structures, defined as $\rho_{\text{fixed}}(\mathbf{x}|c = 1) = \sum_m \alpha_m \delta(\mathbf{x} - \mathbf{x_m})$. It is important to note that while we can still sample from $\rho_{\text{fixed}}$, these samples only represent the cluster centers and lack the local variability inherent in the GMM.

For the experimental datasets where the monomer is the most populous species ($[Mg^{2+}] < 5$ mM), we randomly assigned a holdout set $Y$ consisting of 3000 images for each experimental condition, assuming the holdout images are independent and identically distributed. We sample J=50 times, compute the image-to-structure likelihoods using cryoLike [18], since a full cryoSBI training is computationally more expensive. To robustly estimate the statistical uncertainty of our evaluation, we perform bootstrapping where we resample this the likelihood matrix with replacement along both the image axis and the conformation axis. In practice, to ensure numerical stability when dealing with vanishingly small probabilities in cryo-EM, we compute Eq. 9 in log-space using the logsumexp function. We perform 50 bootstrap iterations for each sampled ensemble. Across the 5 independent mixture models inferred for each condition, we compute the $\text{LLR}(Y_{\text{test}}, \mathbf{x}^{\text{GMM}}, \mathbf{x}^{\text{fixed}})$, this yields a total of 250 evaluations for each condition. This extensive sampling enables us to cross-validate and score ensembles across model conditions.

While the GMM ensemble expands upon fixed structural representations by incorporating covariance parameters, our held-out evaluation framework ensures that any measured advantage reflects genuine predictive power rather than parameter count. Evaluating predictive performance strictly on independent, held-out test images ($Y_{\text{test}}$) inherently controls for parameter-driven bias. Complexity penalties—such as Akaike or Bayesian Information Criteria (AIC/BIC)—are designed to penalize in-sample likelihoods to estimate out-of-sample error; when generalization error is measured directly on held-out data, such penalties are redundant. Furthermore, the GMM covariance parameters do not represent free parameters tuned against cryo-EM image noise, as they were estimated independently from the prior ensemble landscape. Consequently, extra covariance parameters cannot artificially inflate held-out image likelihoods unless they accurately reflect the physical variance of the unseen data.

#### 4 Supplementary Figures

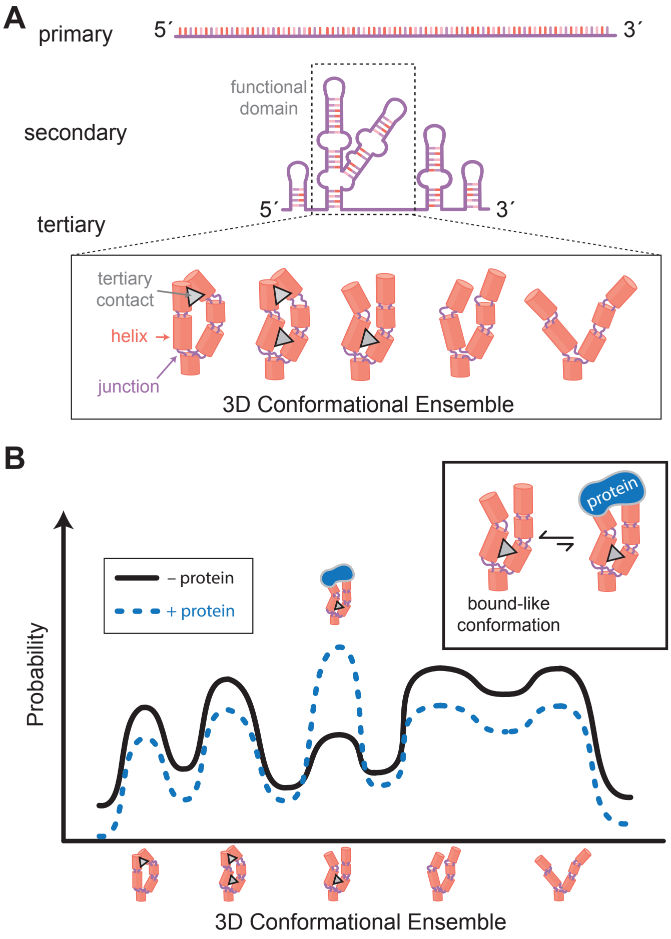

**Supplementary Fig. 1 Functional folded RNAs exist as dynamic conformational ensembles. A** Many RNAs form stable secondary structures but exhibit dynamic tertiary contacts, yielding a conformational ensemble sharing a secondary structure. **B** The ensemble distribution defines the probability of the system adopting specific conformations. This distribution dictates biological function and can be modulated by mutations, binding partners (e.g., protein, shown in blue), and environmental factors. The primary objective of this work is to infer both the conformational and compositional ensemble distributions directly from individual cryo-EM images.

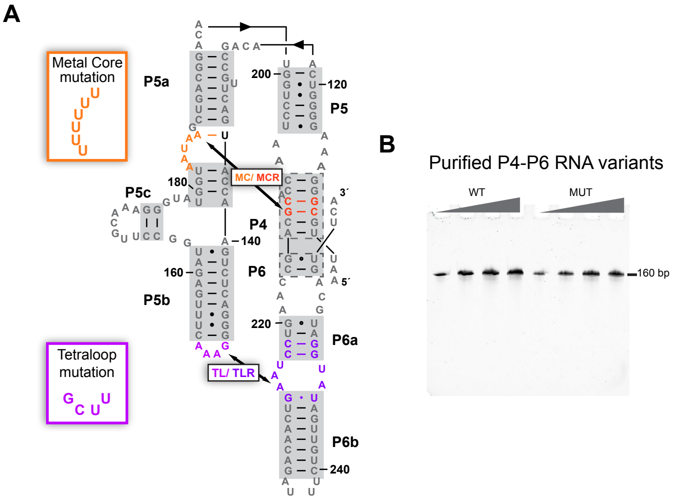

**Supplementary Fig. 2** **A.** Secondary Structure of the P4-P6 domain, showing the MC/MCR and TL/TLR residue-forming contacts. The gray boxes represent the secondary structure elements enforced as constraints in the prior modeling. These blocks represent modular A-form helical segments that function as structural scaffolds connected by junctions, loops, and tertiary interactions, reflecting the hierarchical organization of RNA folding. To make the mutant, we introduce an all uridine rich mutation at the MC and an UUCG mutation at the TL, which abrogates the tertiary contacts of P4-P6 (orange and purple boxes). **B.** Analysis of purified P4-P6 variants by 8% denaturing PAGE.

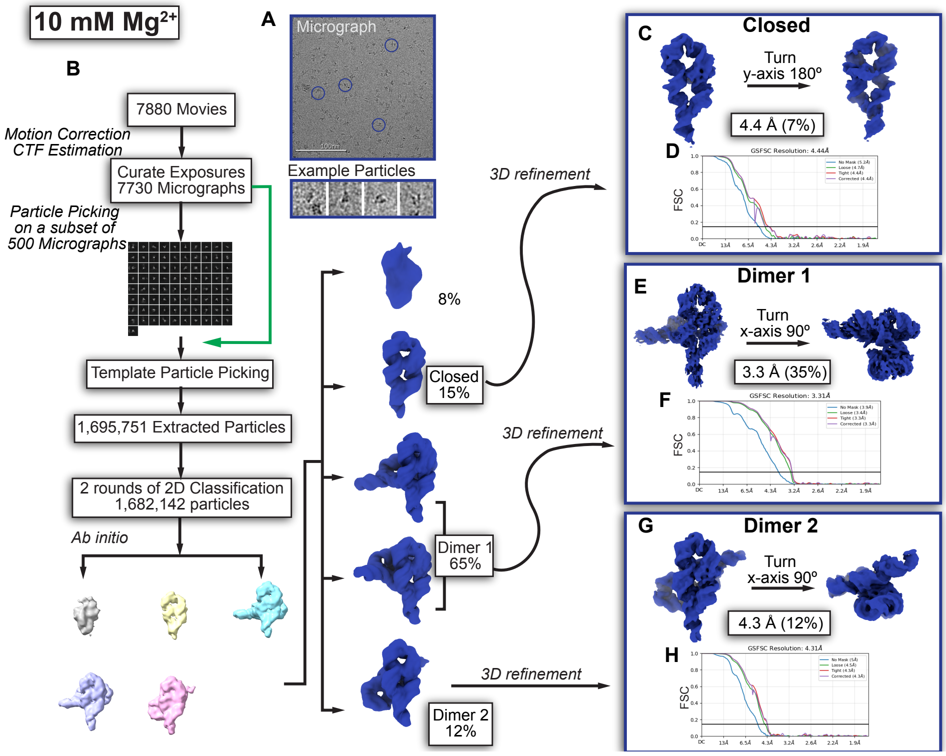

**Supplementary Fig. 3** Cryo-EM data processing workflow for P4-P6 RNA at 10 mM $Mg^{2+}$ **A.** Representative micrograph showing particle distribution on ice, with example particles below. **B.** CryoSPARC workflow for processing P4–P6 RNA datasets collected at 10 mM $Mg^{2+}$, **C.** Final 3D reconstruction map of closed monomeric P4-P6 RNA at 4.4 Åfrom 7 % of the initially extracted particles **D.** Gold-standard Fourier Shell Correlation (FSC) curve at 0.143. **E.** Final 3D reconstruction map of P4-P6 Dimer 1 RNA at 3.3 Å and made with 35 % of the initially extracted particles **F.** Gold-standard FSC curve at 0.143 **G.** Final 3D reconstruction map of P4-P6 Dimer 2 RNA at 4.4 Å and made with 12 % of the initially extracted particles **H.** Gold-standard FSC curve at 0.143.

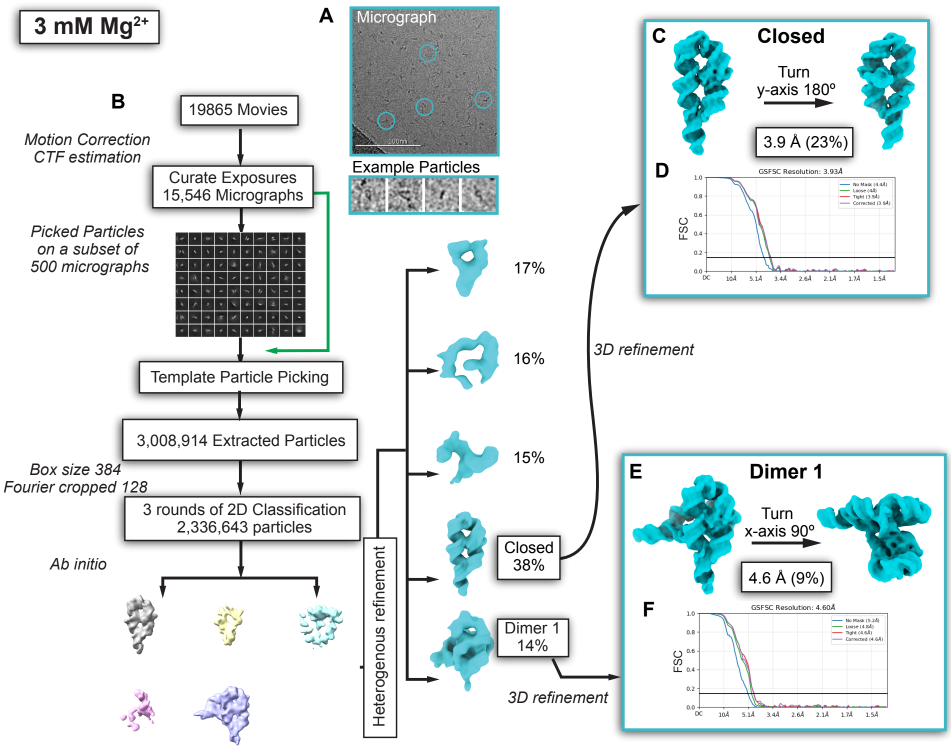

**Supplementary Fig. 4** Cryo-EM data processing workflow for P4-P6 RNA at 3 mM $Mg^{2+}$ **A.** Representative micrograph showing particle distribution on ice, with example particles below. **B.** CryoSPARC workflow for processing P4–P6 RNA datasets collected at 3 mM $Mg^{2+}$, **C.** Final 3D reconstruction map of closed monomeric P4-P6 RNA at 3.9 Å and made with 23 % of the initially extracted particles **D.** Gold-standard FSC curve at 0.143 **E.** Final 3D reconstruction map of P4-P6 Dimer 1 RNA at 4.6 Å and made with 9 % of the initially extracted particles **F.** Gold-standard FSC curve at 0.143.

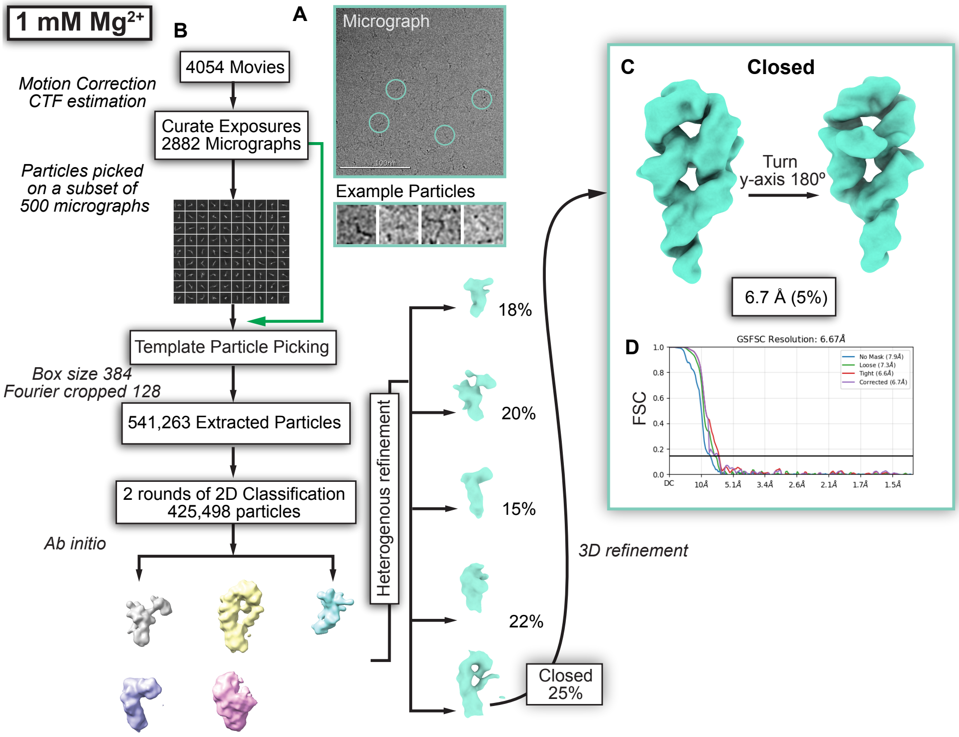

**Supplementary Fig. 5** Cryo-EM data processing workflow for P4-P6 RNA at 1 mM $Mg^{2+}$ **A.** Representative micrograph showing particle distribution on ice, with example particles below. **B.** CryoSPARC workflow for processing P4–P6 RNA datasets collected at 1 mM $Mg^{2+}$, **C.** Final 3D reconstruction map of P4-P6 RNA at 6.7 Å and made with 5 % of the initially extracted particles **D.** Gold-standard FSC curve at 0.143.

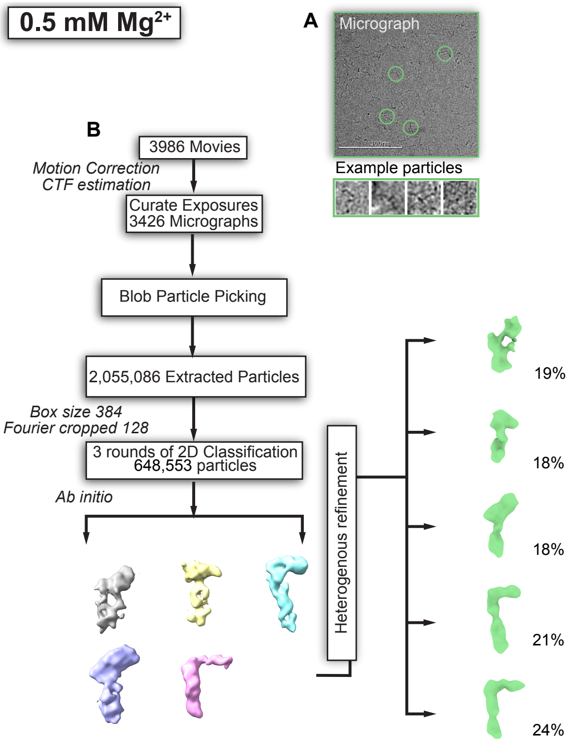

**Supplementary Fig. 6** Cryo-EM data processing workflow for P4-P6 RNA at 0.5 mM $Mg^{2+}$. **A.** Representative micrograph showing particle distribution on ice, with example particles below; **B.** CryoSPARC workflow for processing P4–P6 RNA datasets collected at 0.5 mM $Mg^{2+}$. Pronounced particle heterogeneity under low $Mg^{2+}$ conditions hindered robust particle alignment and refinement, limiting reconstruction of a complete 3D map of the open P4–P6 conformation.

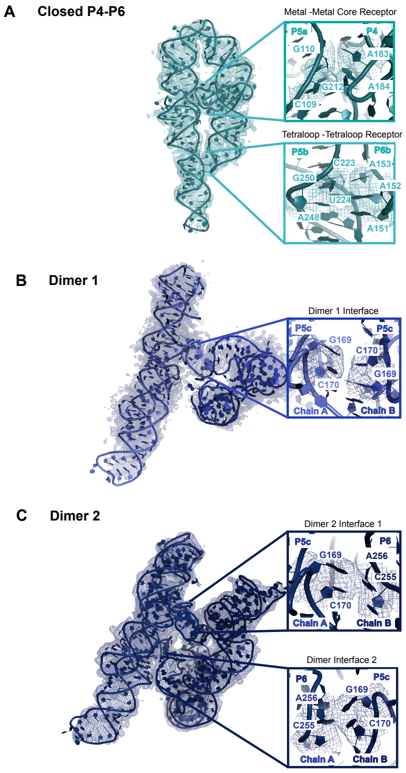

**Supplementary Fig. 7** Structure model building **A.** Closed P4-P6 model docked into a 3.9 Å electron density map. Tertiary contacts shown include the metal core/metal core receptor and tetraloop/tetraloop receptor interactions. **B.** P4-P6 Dimer 1 model docked into a 3.3 Å resolution electron density map showing the dimer 1 interface. **C.** P4-P6 Dimer 2 model docked into a 4.2 Å resolution electron density map showing the two dimer 2 interfaces.

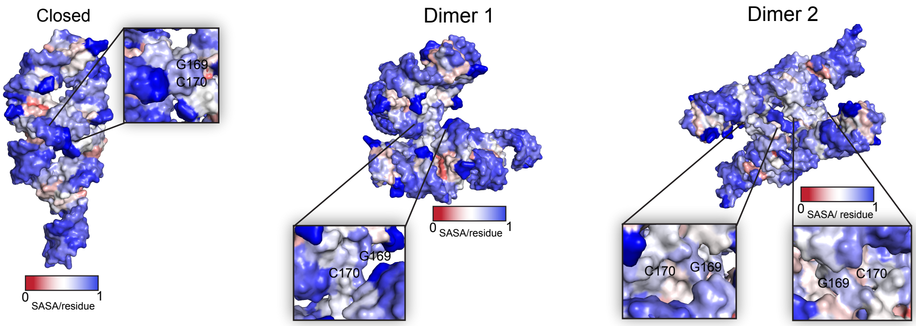

**Supplementary Fig. 8** Solvent accessible surface area (SASA) calculations illustrating progressive burial of the P5 helix (residues G169 and C170) from the closed monomer through Dimer 1 and Dimer 2.

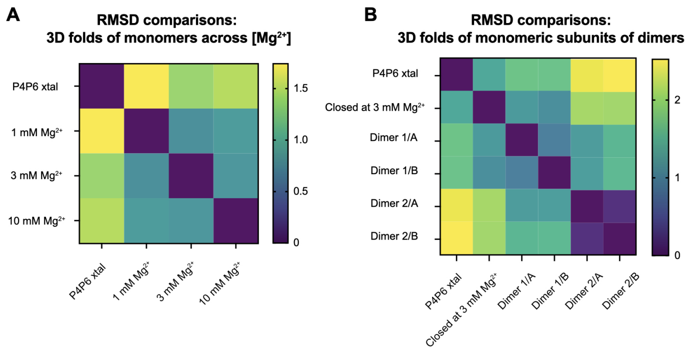

**Supplementary Fig. 9** RMSD comparison of folds across closed monomers for **A**, different [$Mg^{2+}$] and **B**, monomeric subunits in dimers.

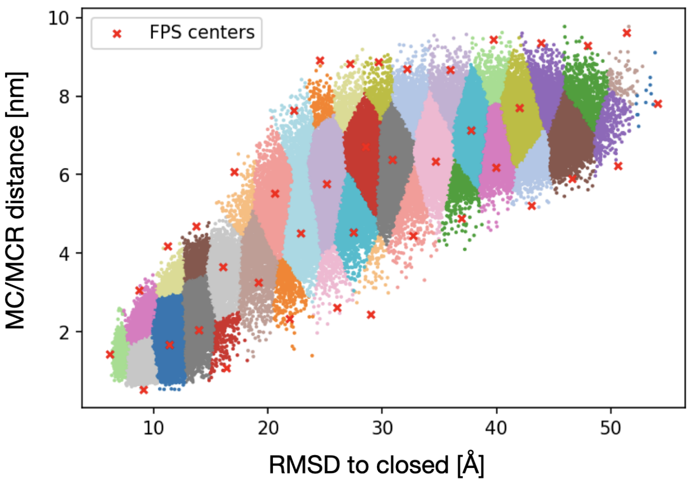

**Supplementary Fig. 10** Example of the prior simulations that span the open-to-closed monomer space, which are tessellated using FPS, as a function of the RMSD to the closed state and MC/MCR distance. FPS centers are shown as red crosses.

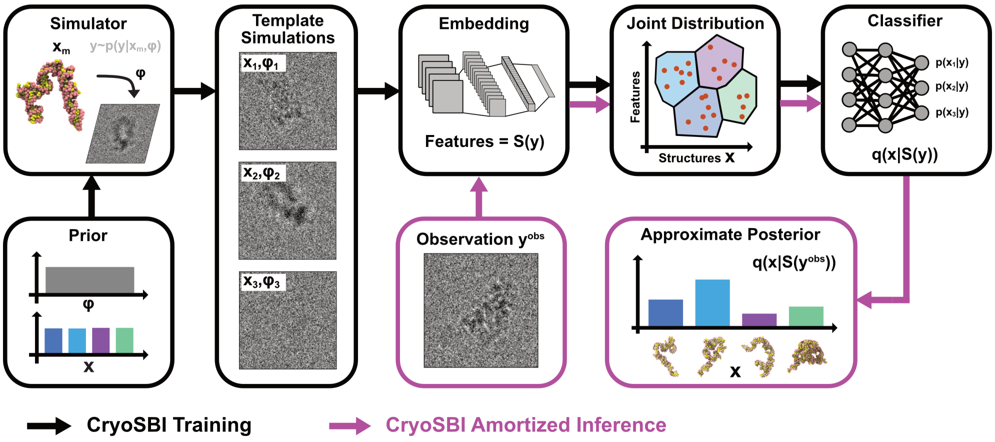

**Supplementary Fig. 11** Schematic of simulation-based inference for cryo-EM over a discrete set of representative structures. Structure labels $\mathbf{x}_m$ are drawn from a uniform categorical prior $p(\mathbf{x}_m)$ over a finite set of representative structures, while nuisance imaging parameters $\boldsymbol{\phi}$, such as rotation, defocus, translation, B-factor, and signal-to-noise ratio, are sampled from a continuous prior $p(\boldsymbol{\phi})$. The labels $\mathbf{x}_m$ and nuisance parameters $\boldsymbol{\phi}$ are then passed to the cryo-EM forward model $p(y)$, which produces a synthetic image $y$. These simulated images are used to jointly train the cryoSBI embedding network and classifier (black boxes): the embedding network featurizes the images as $S(y)$, and the neural classifier learns to assign a probability to each structure label from these features. The trained classifier $q_\psi(\mathbf{x}_m \mid S(y))$ provides a computationally efficient approximation of the structure-image posterior $p(\mathbf{x}_m \mid y)$, taking an image as input and outputting a probability for each representative structure. This enables amortized classification of experimental images $y_i$ against the template set (magenta boxes), returning the full categorical posterior for each particle.

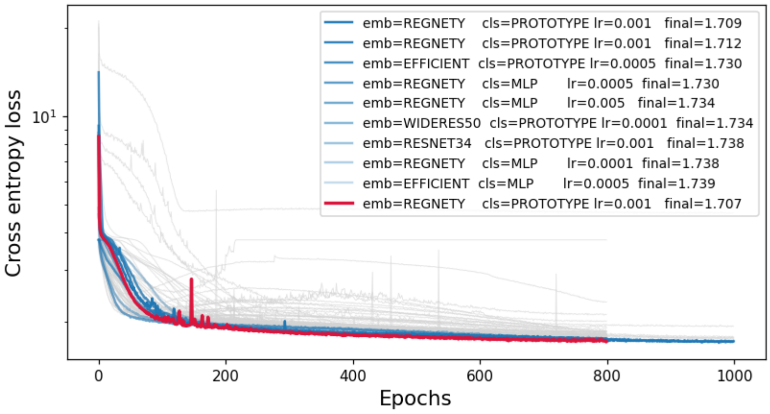

**Supplementary Fig. 12** Hyperparameter scan over 80 training configurations of the cryoSBI classifier. Each curve is the per-epoch training loss of one configuration (gray). The ten configurations with the lowest final loss are highlighted in blue (opacity decreasing with final loss), and the best run is overlaid in red. The lowest-loss configurations consistently combine the RegNetY embedding with the prototype classifier head, supporting this choice for downstream inference.

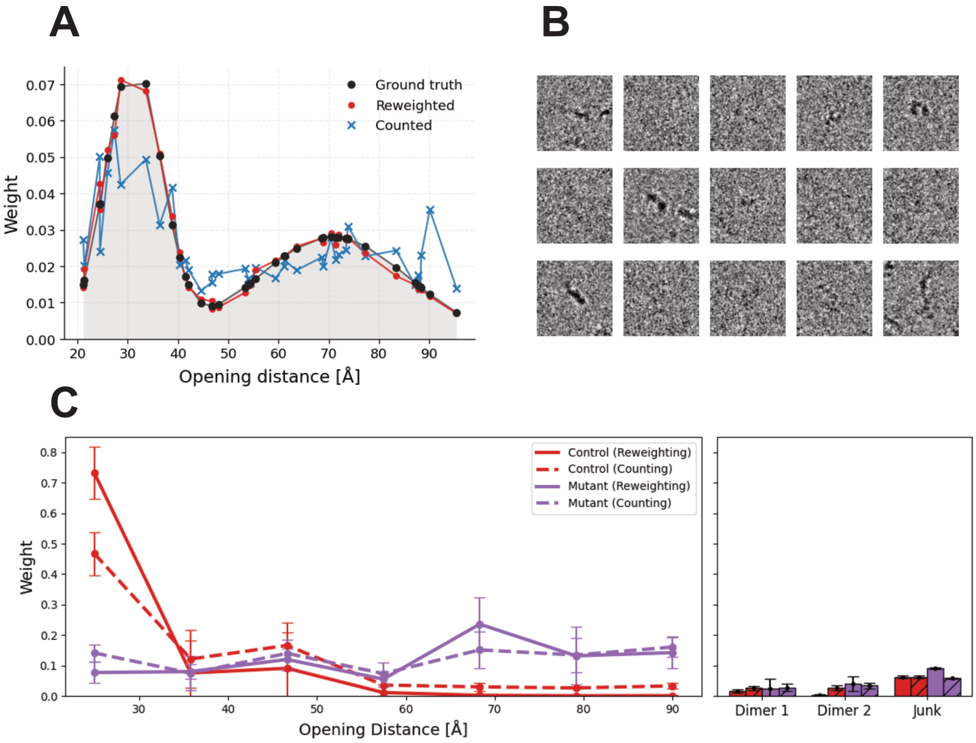

**Supplementary Fig. 13** **A** Recovered density over the opening distance of P4-P6 for $N = 20{,}000$ simulated cryo-EM particles drawn from a bimodal Gaussian target ($\boldsymbol{\mu}_1 = 31.1$ Å, $\boldsymbol{\Sigma}_1 = 5.5$ Å; $\boldsymbol{\mu}_2 = 70.82$ Å, $\boldsymbol{\Sigma}_2 = 15.0$ Å, with equal mixture weights) defined over 40 structural clusters (black dots). The ground-truth target (black) is compared to the distribution obtained from reweighting of the classifier logits (red) and the naive argmax-counting baseline (blue). Reweighting recovers the bimodal target, whereas counting washed out the probability. **B** Representative simulated particles after Gaussian low-pass filtering and per-image normalisation, illustrating the SNR regime (SNR $\in [10^{-4}, 0.1]$) used in panel A. **C** Application to estimate the to experimental cryo-EM distribution for the control and mutant datasets, comparing the population estimates with reweighting with CryoSBI (solid lines, filled bars) and counting with CryoSBI (dashed lines, hatched bars). Left: monomer weights binned by opening distance. Right: weights of the two dimer states and the junk class. Curves and markers show the mean across five independents sets of structural representatives, error bars the standard deviation.

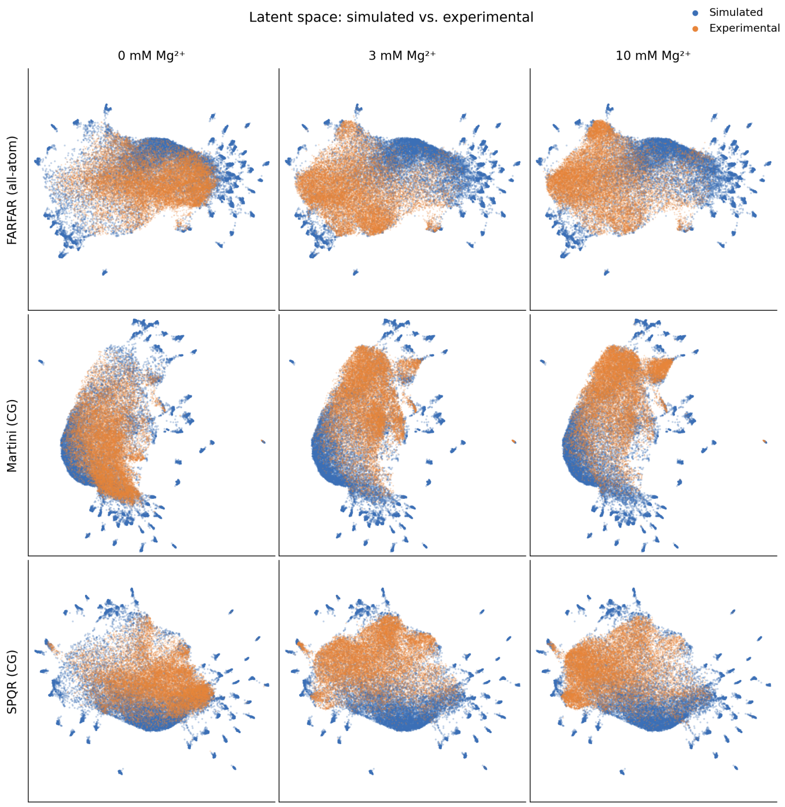

**Supplementary Fig. 14** UMAP projection of the cryoSBI latent space for simulated and experimental particles across prior models and $Mg^{+2}$ conditions. Two-dimensional UMAP projection of the estimator's latent embeddings of simulated particles drawn from the training distribution (blue) and experimental particles (orange); 20,000 particles each. Rows correspond to the three prior ensembles used to train the estimator — FARFAR, Martini and SPQR — and columns to three experimental datasets at increasing $Mg^{2+}$ concentration: 0 mM, 3 mM and 10 mM. For each prior ensemble (row), a single UMAP was fit jointly on the simulated embeddings and all three experimental embeddings, so the three panels within a row share a common projection and are directly comparable.

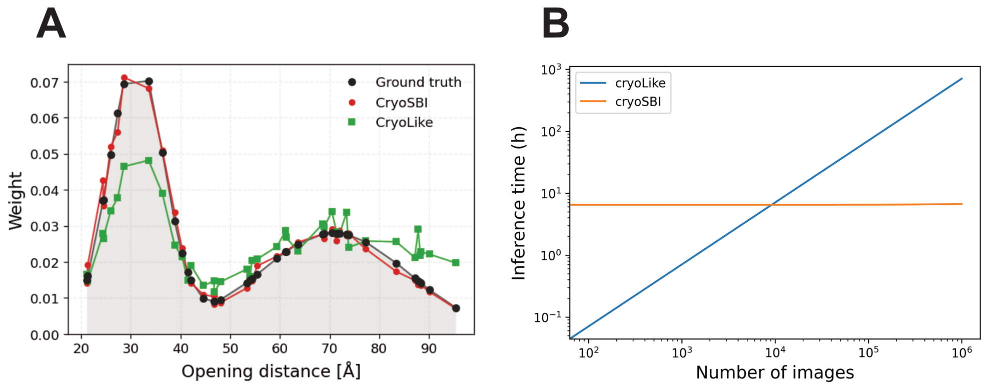

**Supplementary Fig. 15** **A** Recovered density over the opening distance of P4-P6 for $N = 20{,}000$ simulated cryo-EM particles drawn from a bimodal Gaussian target ($\boldsymbol{\mu}_1 = 31.1\,\text{Å}$, $\boldsymbol{\Sigma}_1 = 5.5\,\text{Å}$; $\boldsymbol{\mu}_2 = 70.82\,\text{Å}$, $\boldsymbol{\Sigma}_2 = 15.0\,\text{Å}$, with equal mixture weights) defined over 40 structural clusters (black dots). The ground-truth target (black) is compared to the distribution obtained from reweighting of the classifier logits (red) and from reweighting of likelihood from cryoLike (green). **B** Inference time as a function of the number of images. Once trained, CryoSBI scales very efficiently at roughly 0.7 s per 1000 images, with its training cost needed only once and amortized across all subsequent analyses. CryoLike instead evaluates the likelihood directly, requiring no training but costing around 40 min per 1000 images.

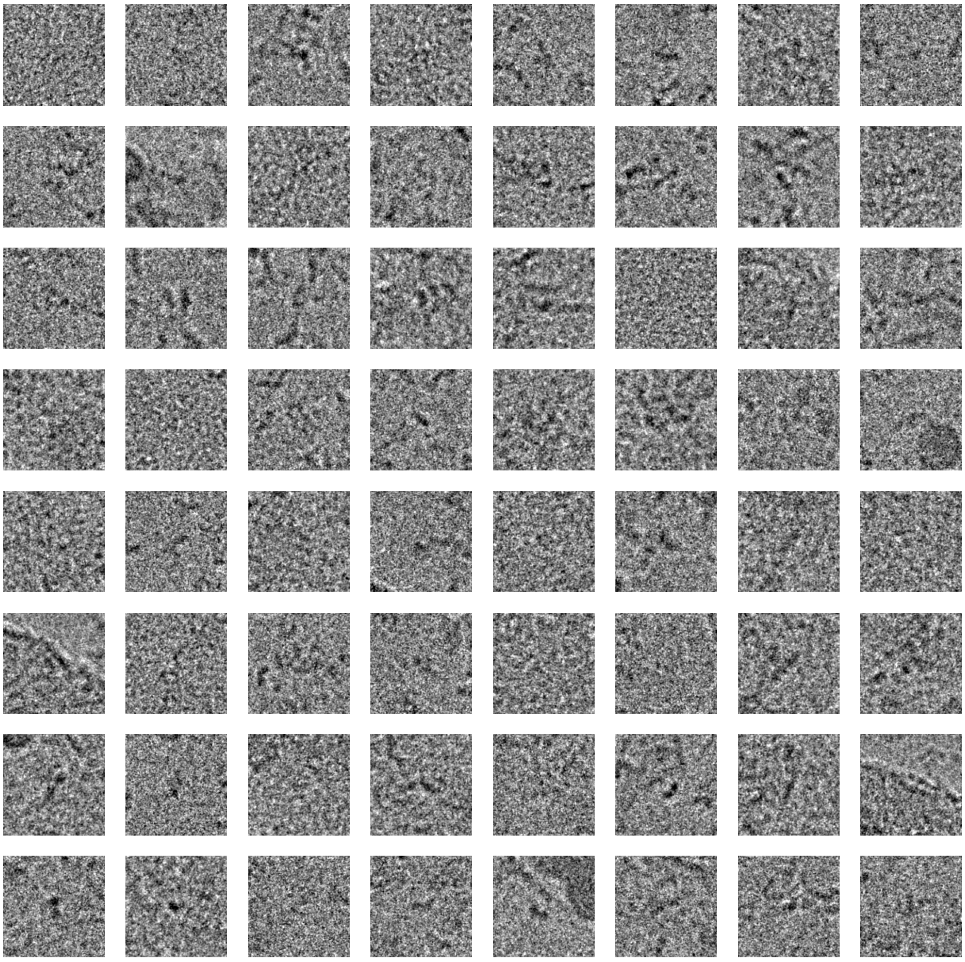

**Supplementary Fig. 16** Random sample of particles assigned to the junk class in the 5 mM $Mg^{+2}$ dataset. Class labels were obtained as the argmax over the classifier logits. The classifier was trained using the martini coarse grained models with 40 structural representatives for the monomers and two dimers.

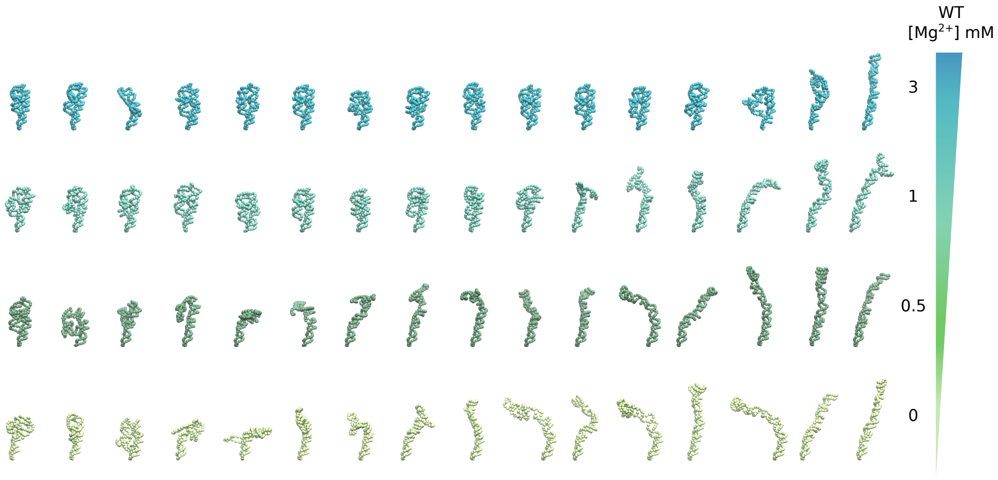

**Supplementary Fig. 17 Conformations sampled from the reweighted monomer GMM ensembles across the $Mg^{2+}$ titration.** Sixteen conformations drawn from the GMM inferred at each $Mg^{2+}$ concentration for wild-type P4–P6 (rows, top to bottom: 3, 1, 0.5 and 0 mM), shown as one CG bead per residue and coloured by condition. Within each row, conformations are ordered by TL/TLR distance, from the most compact on the left to the most extended on the right.

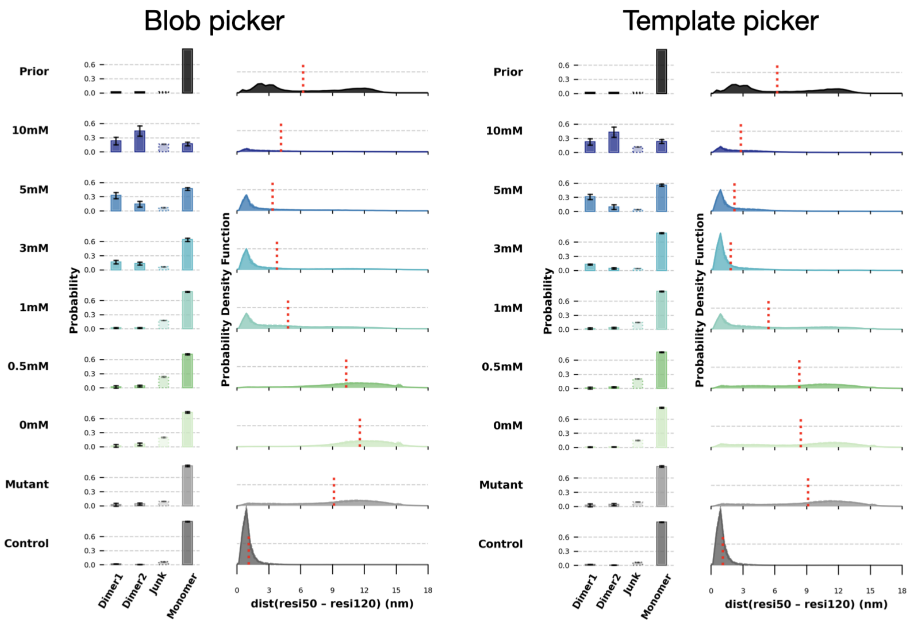

**Supplementary Fig. 18** RNA cryo-EM ensemble reweighting for blob vs template pickers for the different $Mg^{2+}$ concentration datasets and controls. Blob picker has more junk and slightly more weight around the open state for all $Mg^{2+}$ concentrations.

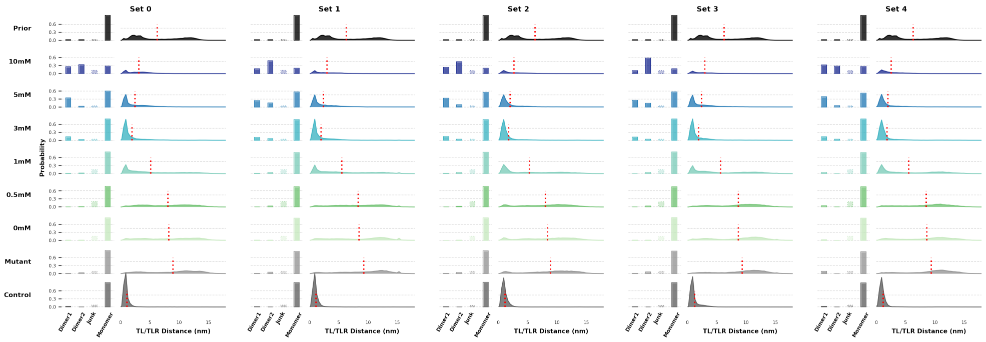

**Supplementary Fig. 19** RNA cryo-EM ensemble reweighting qualitative comparison across independent clustering seeds for the different $Mg^{2+}$ concentration datasets and controls. Qualitative evaluation of five sets with representative FPS cluster centers chosen via different random seeds. Mean and standard deviation values represent the statistical reweighting across the entire ensemble, which are used in all other ensemble reweighting figures.

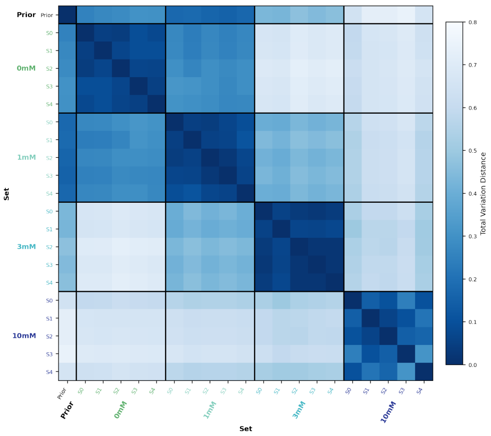

**Supplementary Fig. 20** RNA cryo-EM ensemble reweighting quantitative comparison using the TVD across independent clustering seeds for the 0, 1, 3 and 10 $Mg^{2+}$ concentration datasets. Results between sets are very similar while inter-concentration distributions follow the same trend as in Main Text Figure 3D.

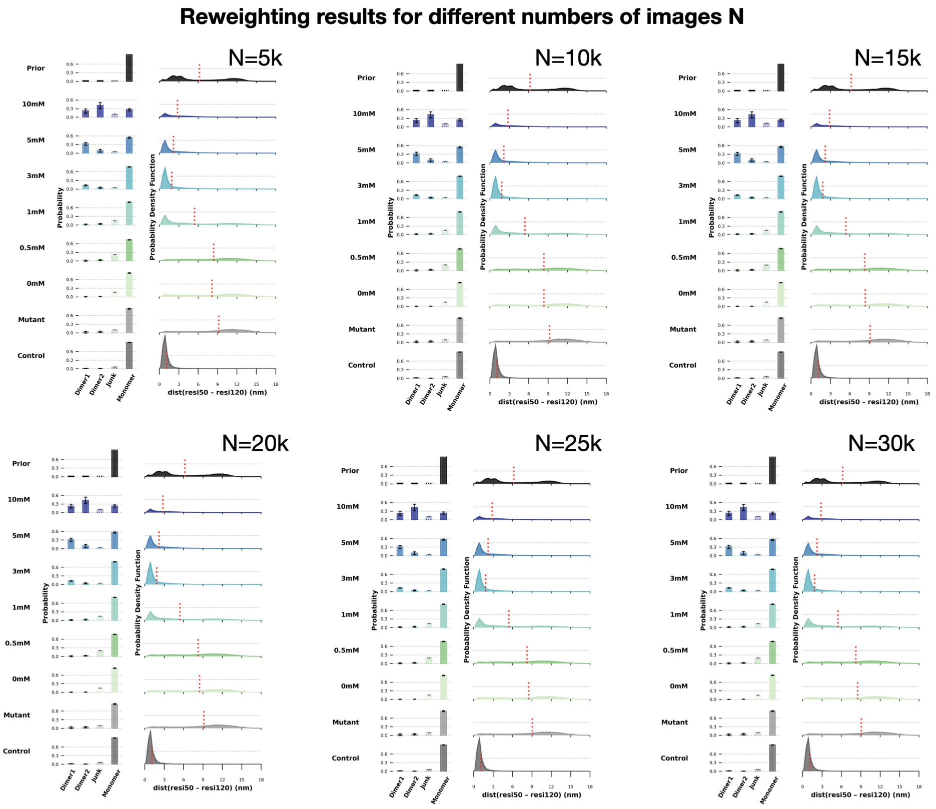

**Supplementary Fig. 21** RNA cryo-EM ensemble reweighting for different number of particles for the different $Mg^{2+}$ concentration datasets and controls. Results are stable after $N \geq 10000$.

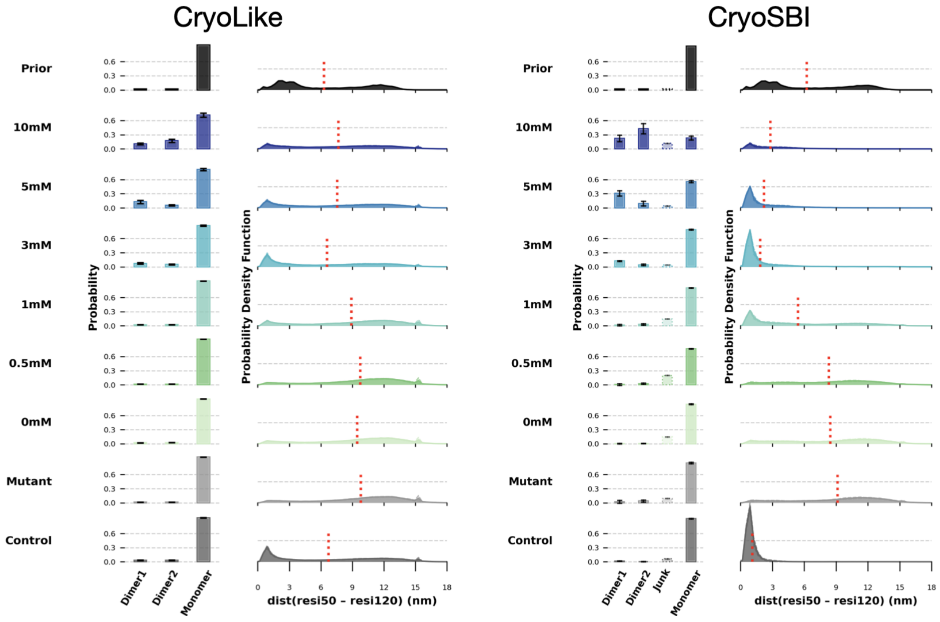

**Supplementary Fig. 22** RNA cryo-EM ensemble reweighting for cryolike vs cryoSBI likelihood calculators for the different $Mg^{2+}$ concentration datasets and controls. Because cryoLike lacks an explicit junk class, the weight from junk particles is likely spread roughly uniformly across all models rather than collected in one class. We hypothesize that this contributes to the flatter distribution inferred by cryoLike, which may also reflect the need for a finer pose grid search, as shown for the synthetic example in Supplementary Figure 15.

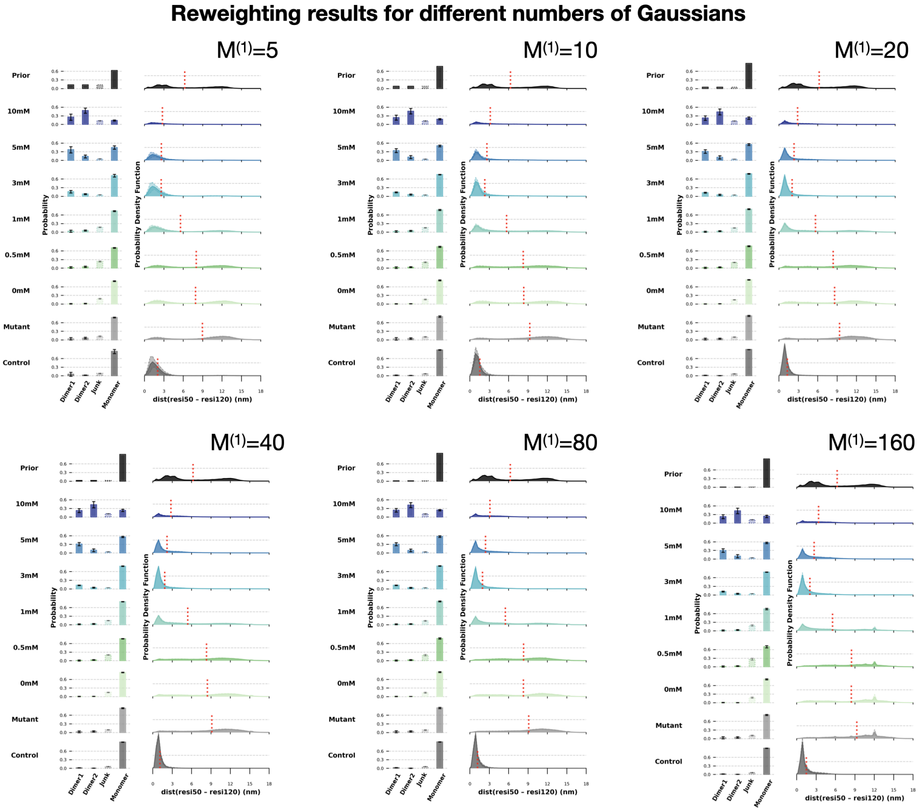

**Supplementary Fig. 23** RNA cryo-EM ensemble reweighting for different number of Gaussians in the monomer GMM ($M^{(1)}$) for the different $Mg^{2+}$ concentration datasets and controls. Note that the total number of models is $M = M^{(1)} + 3$ accounting for the two dimers and junk classes. Inference instabilities, shown by spikyness in the distributions, are found for large $G \geq 80$. This behavior arises because structurally similar conformations can generate statistically indistinguishable images, rendering their corresponding weights unidentifiable [26].

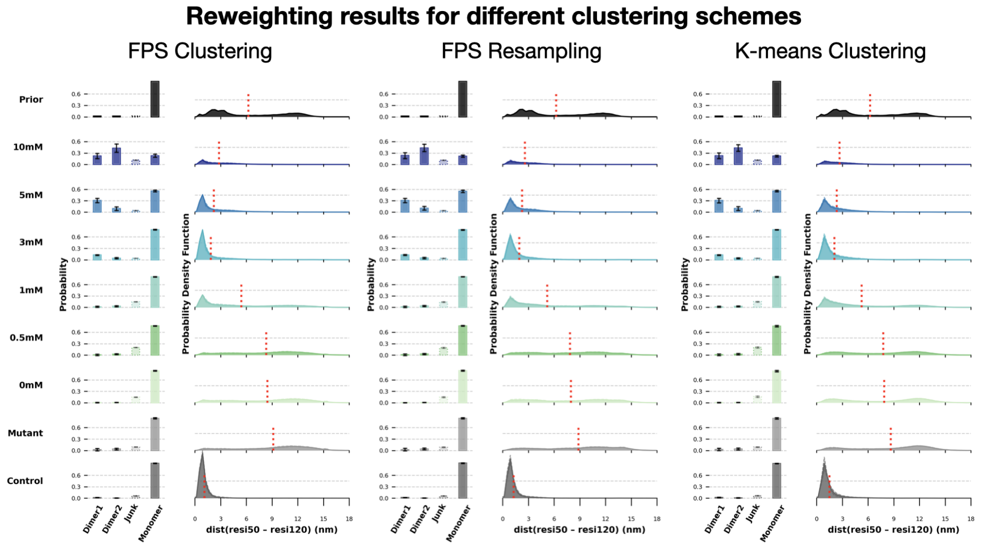

**Supplementary Fig. 24** RNA cryo-EM ensemble reweighting across varying clustering schemes. Results are shown for datasets across different $Mg^{2+}$ concentrations alongside their corresponding controls.

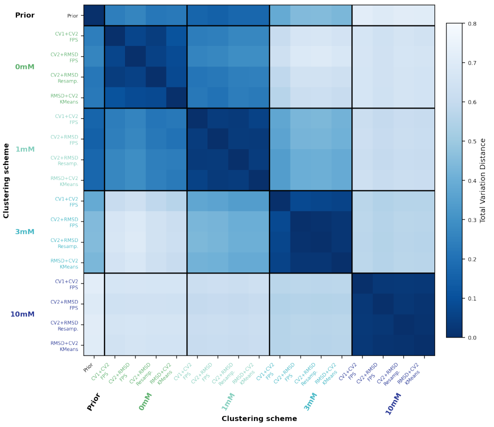

**Supplementary Fig. 25** TVD metric for the ensemble reweighting for clustering along different CVs and schemes, for 0, 1, 3 and 10 $Mg^{2+}$ concentration datasets.

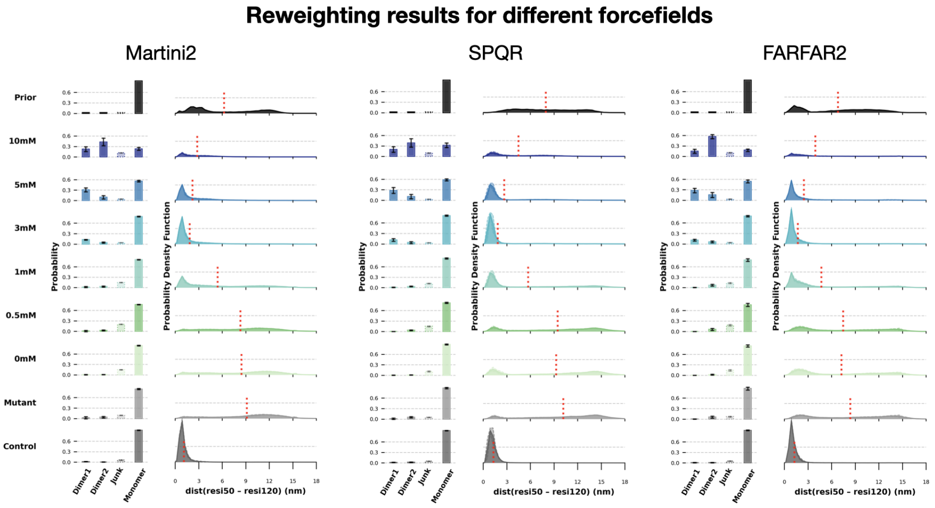

**Supplementary Fig. 26** RNA cryo-EM ensemble reweighting across different force fields and simulators: Martini2, SPAR, and FARFAR2. Results are shown for datasets across different $Mg^{2+}$ concentrations alongside their corresponding controls.

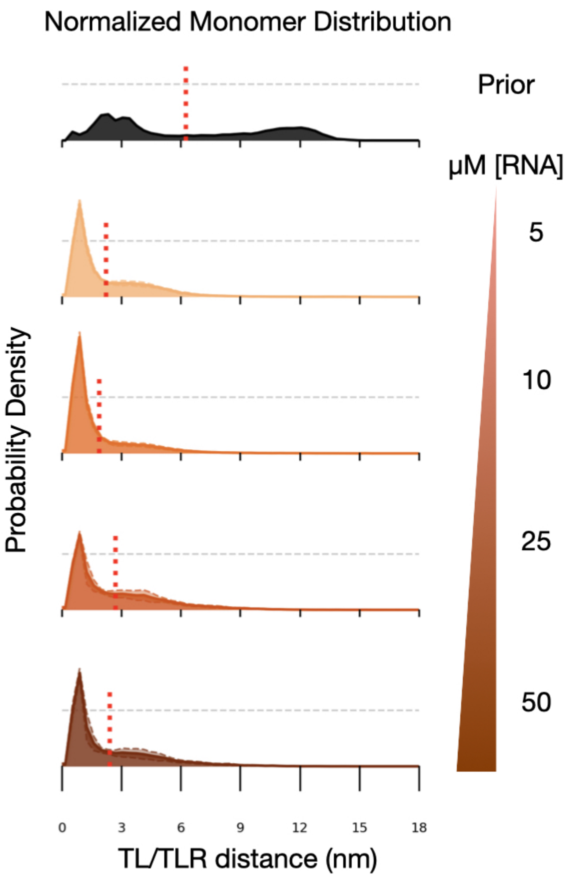

**Supplementary Fig. 27** The inferred monomer conformational distribution is independent of RNA concentration. Normalized probability density over the monomer ensemble projected onto the TL/TLR distance, shown for the prior ensemble (black, top) and for reweighted ensembles obtained at RNA concentrations of 5, 10, 25 and 50 $\mu$M (orange to brown, top to bottom) at 5 mM $Mg^{2+}$. Each distribution is normalized to unit area over the TL/TLR coordinate, so the panels report the shape of the monomer ensemble rather than its absolute population.

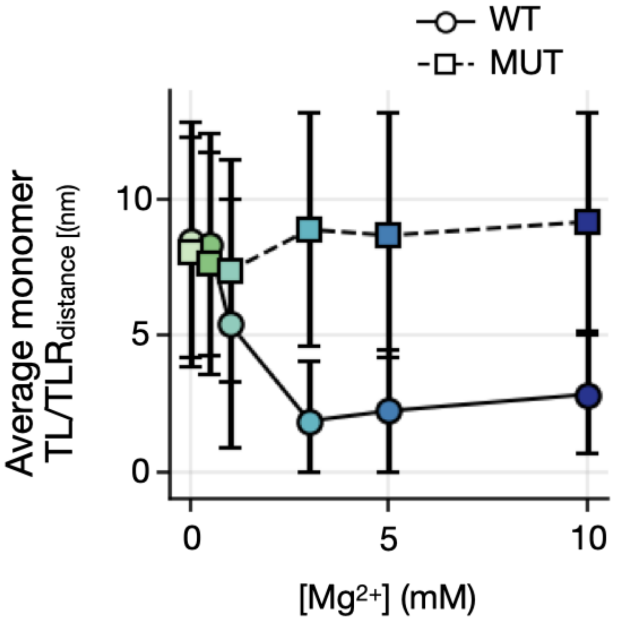

**Supplementary Fig. 28** Averaged monomer TL/TLR$_{\text{distance}}$ as a function of [Mg$^{2+}$] at a constant [RNA] of 50 $\mu$M, for the WT (circle) and MUT (square) titration series. The MUT has been engineered to populate open conformations regardless of [Mg$^{2+}$] conditions.

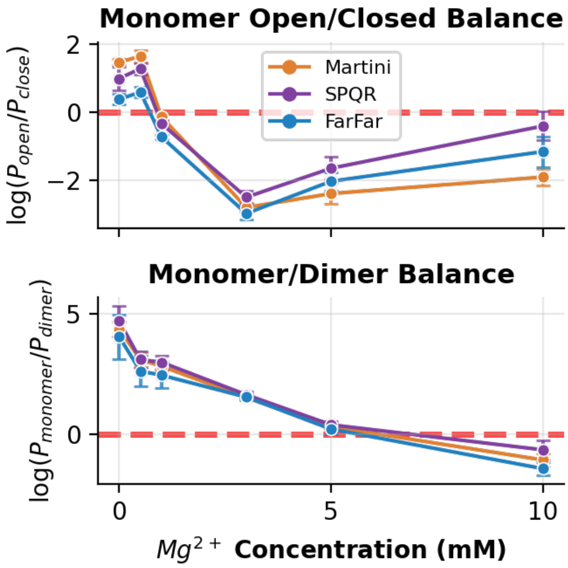

**Supplementary Fig. 29** $Mg^{2+}$-dependent conformational and compositional equilibria of P4–P6 inferred by cryo-EM ensemble reweighting. **(Top)** Log ratio of open to closed monomer populations, $\log(P_{\text{open}}/P_{\text{closed}})$, as a function of $Mg^{2+}$ concentration. Conformations with a TL/TLR distance $\leq$ 3 nm were classified as closed and all others as open. **(Bottom)** Log ratio of monomer to dimer populations, $\log(P_{\text{monomer}}/P_{\text{dimer}})$, over the same titration; dimer weights are the sum of Dimer 1 and Dimer 2. In both panels, colours denote the tool used to generate the prior ensemble: Martini2 (orange), SPQR (purple) and FARFAR2 (blue). Dashed red lines mark equal populations of the two species (log ratio = 0), so that crossings indicate the midpoints of the open-to-closed and monomer-to-dimer transitions. Points are means over five statistical reweighting rounds, each seeded from a different clustering of the prior; error bars are s.d. across rounds.

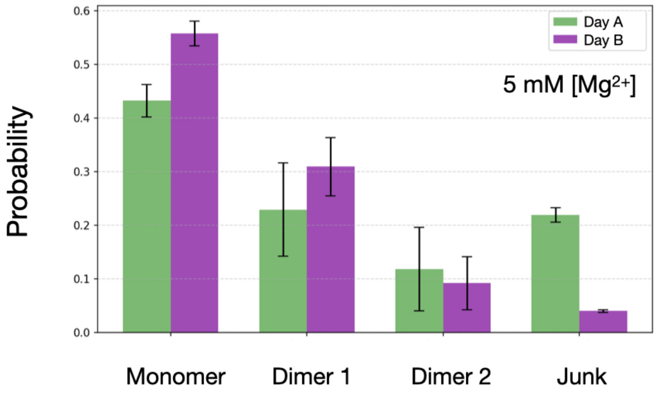

**Supplementary Fig. 30** Inferred weights for samples prepared on different days, folding protocols and microscope sessions, with a constant RNA concentration of 50 $\mu$M and constant $Mg^{2+}$ concentrations of 5 mM. Day A: dataset taken for varying [RNA] and constant [$Mg^{2+}$]. Day B: dataset for ensemble reweighting with constant [RNA] and varying $Mg^{2+}$. Variability was observed within the junk and dimer classes, indicating a higher prevalence of aggregates or degraded particles on specific days as well as between different folding protocols.

### 5 Supplementary Tables

**Table 1** Cryo-EM data collection, processing, 3D reconstructions and model refinement statistics for P4-P6 RNA at 10, 3, 1 and 0.5 mM $Mg^{2+}$

| | P4-P6 RNA | | | |
|---|---|---|---|---|
| | 10 mM $Mg^{2+}$ | 3 mM $Mg^{2+}$ | 1 mM $Mg^{2+}$ | 0.5 mM $Mg^{2+}$ |
| PDB ID | | | | |
| EMDB ID | | | | |
| **Data collection and processing** | | | | |
| Microscope | Titan Krios | | | |
| Voltage (keV) | 300 | | | |
| Camera | Gatan K3 | | | |
| Magnification | 105000 | 105000 | 105000 | 105000 |
| Collection mode | Collection | Super resolution | Super resolution | Super resolution |
| Pixel size at the detector (Å/pixel) | 0.844 | 0.669 | 0.669 | 0.669 |
| Total electron exposure ($e^-/Å^2$) | 62.76 | 55.17 | 54.90 | 54.90 |
| No. of frames | 50 | 31 | 41 | 41 |
| Defocus (µm) | -0.5 – -2.5 | | | |
| Total no. of micrographs | 7,880 | 19,865 | 4,054 | 3,986 |
| Total no. of micrographs used | 7,730 | 15,546 | 2,882 | 3426 |
| Automation software | Leginon | | | |
| **Reconstruction** | | | | |
| Final no. of particles | 117,267, 589,306, 197,620 | 536,353, 206,789 | 21,639 | – |
| Average map resolution (Å) (FSC threshold = 0.143) | 4.4, 3.3, 4.2 | 3.9, 4.6 | 6.7 | – |
| **Model Refinement** | | | | |
| Refinement Software | Phenix | | | – |
| Real or Reciprocal space | Real | | | – |
| Initial Model Used (PDB code) | 1GID | | | – |
| Model resolution (Å) | 4.5, 3.5, 4.3 | 4.1 | 6.8 | – |
| Map sharpening B-factor ($Å^2$) | 183.9, 205.8, 96.2 | 213.2 | 429.6 | – |
| Model Composition, no. of atoms | 3379, 6758, 6758 | 3379 | 3379 | – |
| RNA residues | 158, 316, 316 | 158 | 158 | – |
| Mean B factors ($Å^2$) | 68.87, 75.57, 127.52 | 94.75 | 181.67 | – |
| Model to map scores | | | | |
| $CC_{box}$ | 0.82, 0.67, 0.84 | 0.77 | 0.91 | – |
| $CC_{mask}$ | 0.76, 0.75, 0.75 | 0.76 | 0.81 | – |
| $CC_{volume}$ | 0.74, 0.72, 0.73 | 0.63 | 0.74 | – |
| $CC_{peaks}$ | 0.65, 0.57, 0.64 | 0.75 | 0.79 | – |
| RMSD from ideal values | | | | |
| Bond lengths (Å) | 0.003, 0.003, 0.002 | 0.002 | 0.002 | – |
| Bond angles (°) | 0.612, 0.595, 0.582 | 0.537 | 0.561 | – |
| **Validation** | | | | |
| Molprobity score | 2.72 | 2.74 | 2.97 | – |
| Clashscore, all atoms | 11.31 | 12 | 21.35 | – |

**Table 2** Cryo-EM data collection and processing for $Mg^{2+}$- titration on RNA ensemble reweighting

| | *WT* P4-P6 RNA | | | | | |
|---|---|---|---|---|---|---|
| | 10 mM $Mg^{2+}$ | 5 mM $Mg^{2+}$ | 3 mM $Mg^{2+}$ | 1 mM $Mg^{2+}$ | 0.5 mM $Mg^{2+}$ | 0 mM $Mg^{2+}$ |
| **Data collection** | | | | | | |
| Microscope | Titan Krios | | | | | |
| Voltage (keV) | 300 | | | | | |
| Camera | Gatan K3 | | | | | |
| Magnification | 105000 | | | | | |
| Collection mode | Super-resolution | | | | | |
| Pixel size at the detector (Å/pixel) | 0.669 | | | | | |
| Total electron exposure ($e^-/Å^2$) | 54.68 | | | | | |
| No. of frames | 41 | | | | | |
| Defocus (µm) | -0.5 – -2.5 | | | | | |
| Automation software | Leginon | | | | | |
| Total no. of micrographs | 2373 | 2511 | 1481 | 1659 | 1458 | 2500 |
| **Data Processing** | | | | | | |
| Total no. of micrographs used | 500 | | | | | |
| Particle picking | Template | | | | | |
| Extraction box size (pixels) | 384, fourier cropped to 128 | | | | | |
| Total no. of particles | 34,611 | 61,387 | 44,577 | 73,416 | 81,160 | 64,792 |

| | *MUT* P4-P6 RNA | | | | | |
|---|---|---|---|---|---|---|
| | 10 mM $Mg^{2+}$ | 5 mM $Mg^{2+}$ | 3 mM $Mg^{2+}$ | 1 mM $Mg^{2+}$ | 0.5 mM $Mg^{2+}$ | 0 mM $Mg^{2+}$ |
| **Data collection** | | | | | | |
| Microscope | Titan Krios | | | | | |
| Voltage (keV) | 300 | | | | | |
| Camera | Gatan K3 | | | | | |
| Magnification | 105000 | | | | | |
| Collection mode | Super-resolution | | | | | |
| Pixel size at the detector (Å/pixel) | 0.669 | | | | | |
| Total electron exposure ($e^-/Å^2$) | | 54.96 | | | 55.04 | |
| No. of frames | 41 | | | | | |
| Defocus (µm) | -0.5 – -2.5 | | | | | |
| Automation software | Leginon | | | | | |
| Total no. of micrographs | 3,665 | 3,563 | 3,477 | 3,213 | 3,105 | 3,226 |
| **Data Processing** | | | | | | |
| Total no. of micrographs used | 500 | | | | | |
| Particle picking | Template | | | | | |
| Extraction box size (pixels) | 384, fourier cropped to 128 | | | | | |
| Total no. of particles | 54,481 | 55,425 | 69,452 | 25,591 | 25,238 | 37,604 |

**Table 3** Cryo-EM data collection and processing for P4-P6 RNA concentration titration on RNA ensemble reweighting

| | *WT* P4-P6 RNA | | | |
|---|---|---|---|---|
| | 5 $\mu$M | 10 $\mu$M | 25 $\mu$M | 50 $\mu$M |
| **Data collection** | | | | |
| Microscope | Titan Krios | | | |
| Voltage (keV) | 300 | | | |
| Camera | Gatan K3 | | | |
| Magnification | 105000 | | | |
| Collection mode | Super-resolution | | | |
| Pixel size at the detector (Å/pixel) | 0.669 | | | |
| Total electron exposure ($e^-$/Å$^2$) | 54.92 | | | |
| No. of frames | 41 | | | |
| Defocus (µm) | -0.5 – -2.5 | | | |
| Automation software | Leginon | | | |
| Total no. of micrographs | 3,836 | 3,601 | 3,963 | 3,318 |
| **Data Processing** | | | | |
| Total no. of micrographs used | 500 | | | |
| Particle picking | Template | | | |
| Extraction box size (pixels) | 384, fourier cropped to 128 | | | |
| Total no. of particles | 44,192 | 53,268 | 83,776 | 116,229 |

**Table 4** List of P4-P6 RNA constructs and PCR Primers. ***T7 promoter sequence**

| | P4-P6 RNA Constructs |
|---|---|
| *WT* | GCG**AGTGAATTCTAATACGACTCACTATA**<br>**GG**AATTGCGGGAAAGGGGTCAACAGCCGT<br>TCAGTACCAAGTCTCAGGGGAAACTTTGA<br>GATGGCCTTGCAAAGGGTATGGTAATAAGCTG<br>ACGGACATGGTCCTAACCACGCAGCCAAGTC<br>CTAAGTCAACAGATCTTCTGTTGATATGGAT<br>GCAGTTCA |
| *MUT* | GCG**AGTGAATTCTAATACGACTCACTATA**<br>**GG**AATTGCGGGAAAGGGGTCAACAGCCGT<br>TCAGTACCAAGTCTCAGGG**ttcg**CTTTGA<br>GATGGCCTTGCAAAGGGTATGGT**tttttt**GCTG<br>ACGGACATGGTCCTAACCACGCAGCCAAGTC<br>CTAAGTCAACAGATCTTCTGTTGATATGGAT<br>GCAGTTCA |
| | PCR primers |
| Forward Primer | AGTGAATTCTAATACGACTCACTATAGG |
| Reverse Primer | TGAACTGCATCCATATCAACAG |

**Table 5** CryoSBI training wall-clock times on a single A100 GPU, averaged over the five independent replicates

| Model | Training time (h) |
|---|---|
| *Single-bead representation* | |
| FARFAR, 40 Clusters | 4.21 |
| Martini, 40 Clusters | 4.24 |
| SPQR, 40 Clusters | 4.44 |
| *Coarse-grained representation* | |
| SPQR, 40 clusters | 6.28 |
| Martini, 40 clusters | 6.46 |
| Martini, 160 clusters | 6.82 |
| Martini, 40 clusters (k-means) | 6.84 |
| Martini, 40 clusters (CV1–CV2) | 6.88 |
| Martini, 80 clusters | 6.89 |
| Martini, 40 resampling | 6.94 |
| Martini, 20 clusters | 7.03 |
| Martini, 10 clusters | 7.31 |
| Martini, 5 clusters | 7.64 |
| *All-atom representation* | |
| FARFAR, 40 Clusters | 19.64 |
